# Chromatin remodeler Chd7 regulates photoreceptor development and outer segment length

**DOI:** 10.1101/2022.05.30.494019

**Authors:** Laura A. Krueger, Jessica D. Bills, Zun Yi Lim, Jennifer M. Skidmore, Donna M. Martin, Ann C. Morris

## Abstract

Mutations in the chromatin remodeling factor CHD7 are the predominant cause of CHARGE syndrome, a congenital disorder that frequently includes ocular coloboma. Although CHD7 is known to be required for proper ocular morphogenesis, its role in retinal development has not been thoroughly investigated. Given that individuals with CHARGE syndrome can experience visual impairment even in the absence of coloboma, a better understanding of CHD7 function in the retina is needed. In this study, we characterized the expression pattern of Chd7 in the developing zebrafish and mouse retina and documented ocular and retinal phenotypes in *Chd7* loss-of-function mutants. Zebrafish Chd7 was expressed throughout the retinal neuroepithelium when retinal progenitor cells were actively proliferating, and later in subsets of newly post-mitotic retinal cells. At stages of retinal development when most retinal cell types had terminally differentiated, Chd7 expression remained strong in the ganglion cell layer and in some cells in the inner nuclear layer. Intriguingly, strong expression of Chd7 was also observed in the outer nuclear layer where it was co-expressed with markers of post-mitotic cone and rod photoreceptors. Expression of mouse CHD7 displayed a similar pattern, including expression in the ganglion cells, subsets of inner nuclear layer cells, and in the distal outer nuclear layer as late as P15. Two different mutant *chd7* zebrafish lines were characterized for ocular and retinal defects. These mutants displayed microphthalmia, reduced numbers of cone photoreceptors, and truncated rod and cone photoreceptor outer segments. Reduced cone photoreceptor number and abnormal outer segments were also observed in heterozygous *Chd7* mutant mice. Taken together, our results in zebrafish and mouse reveal a conserved, previously undescribed role for Chd7 in retinal development and photoreceptor outer segment morphogenesis. Moreover, our work suggests an avenue of future investigation into the pathogenesis of visual system defects in CHARGE syndrome.

**Highlights:**

- Chd7 is expressed in both retinal progenitor cells and in differentiated retinal neurons, including post-mitotic rod and cone photoreceptors.
- Loss of Chd7 results in a significant decrease in cone photoreceptors in both zebrafish and mouse.
- Cone and rod photoreceptor outer segments are truncated in *chd7* mutants, suggesting a heretofore unappreciated role for Chd7 in outer segment morphogenesis.

## 1. Introduction

Development of the vertebrate visual system requires the precise spatial and temporal coordination of numerous gene regulatory networks and signaling pathways. Early morphogenetic events lead to the formation of the optic cup and the structural architecture of the eye; within the developing retina, this is followed by specification and differentiation of the seven major retinal cell types in a largely conserved spatiotemporal sequence. This complex process culminates in a highly organized neural retina that is capable of capturing light and converting it into an electrical signal that can then be transmitted to the brain and interpreted as an image.

Disruptions in eye development can result in congenital ocular malformations such as microphthalmia, anophthalmia, and coloboma (collectively referred to as MAC), as well as retinal cell defects, and are a significant cause of pediatric blindness (Francis, 2006; Williamson and FitzPatrick, 2014). These ocular abnormalities often occur as a part of larger syndromic disorders. One example is CHARGE syndrome, a genetic neural cristopathy characterized by coloboma, heart defects, choanal atresia, growth retardation, genital abnormalities, and ear abnormalities (Hall, 1979; Hittner et al., 1979; Hsu et al., 2014). Based on clinical reports, coloboma is present in over 80% of clinically diagnosed CHARGE syndrome patients(Russell-Eggitt et al., 1990). However, CHARGE syndrome is also associated with other ocular complications, including microphthalmia, optic nerve hypoplasia, nystagmus, cataracts, amblyopia, microcornea, strabismus and rarely angle closure defects and retinal detachment (Aramaki et al., 2006; Guirgis and Lueder, 2003; Lalani et al., 2006; Natung et al., 2014; Russell-Eggitt et al., 1990; Strömland et al., 2005; A L Tellier et al., 1998; A.L Tellier et al., 1998). A recent study of visual function in 14 CHARGE syndrome patients found that all had some degree of visual impairment, even in the absence of structural malformations such as coloboma (Onesimo et al., 2021). These results suggest that the visual system disruptions associated with CHARGE syndrome may involve retinal developmental defects in addition to problems with ocular morphogenesis.

Decades after the initial description of CHARGE syndrome, pathogenic variants in the chromatin remodeling factor CHD7 were identified as the predominant genetic cause (Jongmans et al., 2006; Vissers et al., 2004; Zentner et al., 2010). CHD7 is a member of the chromodomain helicase-DNA binding domain (CHD) family of proteins, initiating nucleosome remodeling that is essential for transcription at target loci (Manning and Yusufzai, 2017). CHD7 has been shown to work in a tissue-specific manner to control target gene expression, by acting alone or as member of a complex with other transcription factors (Bajpai et al., 2010; Ho and Crabtree, 2010; Schnetz et al., 2009).

CHD7 has also been shown to promote neurogenesis in many of the tissues involved in CHARGE syndrome. Disruption of mouse CHD7 results in defects in the development of specific sensory neurons in the auditory, vestibular, and olfactory systems (Hurd et al., 2010; Layman et al., 2009). These data suggest that CHD7 may also play a role in the development of sensory neurons in the retina, but studies specifically addressing this question have been limited. One study used mouse conditional knockouts to delete CHD7 from various embryonic tissues that contribute to the eye; this work demonstrated that expression of CHD7 is required in the neural ectoderm for proper ocular morphogenesis of the optic cup and optic fissure closure (Gage et al., 2015). Although the authors also noted CHD7 expression in the developing mouse retina, the effects of loss of CHD7 on retinal neurogenesis were not extensively characterized (Gage et al., 2015). General descriptions of microphthalmia, coloboma, and anterior segment defects have been noted in zebrafish models of Chd7 deficiency, but potential retinal phenotypes have not been investigated (Breuer et al., 2020; Cloney et al., 2018; Jacobs-Mcdaniels and Albertson, 2011). One study using morpholino-mediated knockdown in zebrafish observed gross retinal lamination defects upon loss of Chd7, indicating a requirement for Chd7 in retinal organization (Patten et al., 2012). Clearly, more in-depth studies are needed to provide a better understanding of the pathogenetic mechanisms leading to CHD7-associated ocular complications of CHARGE syndrome, and to determine whether these include retinal differentiation defects.

In this study, we analyzed Chd7 expression in the developing retina and investigated retinal phenotypes in zebrafish and mouse *Chd7* mutants. We demonstrate that in the zebrafish and mouse retina, Chd7 is expressed not only in early retinal progenitor cells, but also later in subsets of differentiated retinal neurons, including mature rod and cone photoreceptors. Importantly, we show that loss of Chd7 causes a significant decrease in cone photoreceptor number and truncated photoreceptor outer segments in both species. Taken together, this work suggests a novel, conserved role for Chd7 in retinal development and photoreceptor outer segment morphogenesis.

## 2. Material and Methods

### 2.1 Zebrafish Lines and Maintenance

Zebrafish were bred, raised, and housed at 28.5°C on a 14-hour light:10-hour dark cycle in compliance with established protocols for zebrafish husbandry (Westerfield, 2000). The Tg(3.2TαC:EGFP) transgenic line (TαC:EGFP) has been previously described and was generously provided by Susan Brockerhoff (University of Washington, Seattle, WA) (Kennedy et al., 2007). The Tg(XlRho:EGFP) transgenic line (XOPs:GFP) has been previously described and was obtained from James Fadool (Florida State University, Tallahassee, FL) (Fadool, 2003). The *chd7^sa19732^* mutant line was originally generated by the Zebrafish Mutation Project at the Wellcome Sanger Institute (Kettleborough et al., 2013) and was obtained from the Zebrafish International Resource Center (ZIRC, Eugene, OR). Heterozygous *chd7* adult CRISPR mutants were provided by Jason Berman (Dalhousie University Halifax, Nova Scotia, CA/Children’s Hospital of Eastern Ontario Research Institute, Ottawa, Ontario, CA) (Cloney et al., 2018). Wildtype zebrafish (AB strain) were obtained from the Zebrafish International Resource Center (ZIRC, Eugene, OR). All animal procedures were carried out in accordance with guidelines established by the University of Kentucky Institutional Animal Care and Use Committee and the ARVO Statement for the Use of Animals in Ophthalmic and Vision Research.

### 2.2 Zebrafish Genomic DNA extraction and amplification

Genomic DNA (gDNA) was extracted from whole embryos or tails. Tissue was placed in 50mM sodium hydroxide and incubated at 95°C for digestion. The solution was neutralized with 1 M Tris-HCl, pH 8.0. The gDNA was PCR amplified using primers described in Supplemental Table 1. Products were visualized on 2% agarose gels and successful amplification reactions were prepared for Sanger sequencing with Exo-CIP™ Rapid PCR Cleanup Kit according to the manufacturer’s protocol (New England Biolabs: E1050S, Ipswich, MA). Samples were then sequenced with amplification primers (Eurofins Genomics Services, Louisville, KY).

### 2.3 Mouse Lines and Maintenance

*Chd7^Gt/+^* (Hurd et al., 2007) (JAX stock 030659) mice were maintained by backcrossing B6129SF1/J (JAX stock 101043) mice to N15. Timed pregnancies were established between *Chd7^Gt/+^* male mice and *Chd7^+/+^* female mice from the colony. The day of plug identification was designated as day 0.5. The day of birth was designated as P1. All procedures were approved by The University of Michigan University Institutional Animal Care & Use Committee (IACUC). Ear punches were collected from mice (P14-P21) and tail snips from experimental embryos and pups. DNA was extracted using a HotShot method (Truett et al., 2000) and samples were analyzed by PCR (primer sequences provided in Supplementary Table 1) using cycling parameters designed by the Jackson Laboratory (Protocol 31768).

### 2.4 Mouse Tissue Preparation

Embryos (E15.5) and heads (skin removed E18.5 and P1) were dissected before fixation in 4% PFA for 2h at room temperature. Specimens were washed three times in 1x PBS and transferred to 70% EtOH and dehydrated and embedded in paraffin wax using a Tissue Tek (Torrance, CA) embedding machine. Paraffin embedded tissues were stored at room temperature until sectioning. Eyes were enucleated from pups (P6, P10, P15) as previously described (Garnai et al., 2019). Briefly, pups were anesthetized with CO2 and euthanized by cervical dislocation. Skin around the eyes was snipped with dissecting scissors to expose the eyes. Eyes were enucleated with preservation of the optic nerves using Excelta 7-P1 curved forceps. Eyes were fixed in 4% PFA overnight at 4°C, then washed in 1x PBS and incubated overnight in 10% followed by 30% sucrose at 4°C.

### 2.5 Western Blot

Protein lysate was extracted from a pool of 75 zebrafish heads collected from 3 dpf wildtype, heterozygous, and *chd7* mutant zebrafish larvae in 1x RIPA buffer. Protein was quantified on a BioSpectrometer (Eppendorf) with a Bradford Assay (Bradford Reagent-E530-1L). Protein lysate was diluted with an appropriate amount of 4x Loading dye and incubated at 98°C for 5 minutes. 20μg of each respective sample was loaded onto a 1mm 3-8% Tris-Acetate Mini Protein gel (NuPAGE) along with 10μl of Spectra Multicolor High Range protein ladder (Thermofisher). The gel was run at 80V for 15 minutes and then increased to 120V until the dye front reached the bottom. Wet transfer onto a 0.2 μm nitrocellulose membrane (BioRad) was performed overnight at 4°C with a stir bar and ice pack at 30mA for approximately 24hrs. The blot was then blocked in 5% milk PBS-T for 2.5 hours at room temperature, and incubated with either anti-CHD7 (1:500, Boster Bio DZ01533) or anti-Vinculin (1:2000, Boster Bio PA1781) primary antibody in 0.5% milk PBS-T overnight at 4°C on a shaker. The membrane was washed 3 times for 5 minutes each with PBS-T on a shaker at room temperature. The blot was resuspended in 0.5% milk PBS-T and incubated in anti-rabbit secondary antibody (1:1000, Santa Cruz sc-2357) for one hour at room temperature. After incubation, it was washed again 3 times for 5 minutes each with PBS-T on a shaker at room temperature. The membrane was imaged on an Amersham Imager 680.

### 2.6 Immunohistochemistry

Zebrafish embryos were fixed overnight in 4% paraformaldehyde, then incubated overnight in 10% followed by 30% sucrose at 4°C. 10 μm transverse zebrafish and mouse cryosections were collected on a Leica CM1900 crysostat (Leica Biosystems, Buffalo Grove, IL). 10 μm paraffin mouse sections were collected on a Thermo Scientific Shandon Finesse ME Microtome (Thermo Scientific, Waltham, MA).

Immunohistochemistry was performed on sections as previously described (Forbes-Osborne et al., 2013), using the following primary antibodies: anti-zebrafish Chd7 (rabbit, 1:500; Boster Bio:DZ01533, Pleasanton, CA); anti-mouse CHD7 (rabbit, 1:2500, Cell Signaling Technologies:6505, Danvers, MA); 4C12, which labels zebrafish rod photoreceptors (mouse, 1:100, J. Fadool, FSU, Tallahassee, FL); 1D1, which labels zebrafish rhodopsin (mouse, 1:100, J. Fadool, FSU, Tallahassee, FL); Zpr-1, which labels zebrafish red-green double cones (mouse, 1:20, ZIRC); Peanut agglutinin (PNA)-lectin conjugated to Cy5 (Vector Labs cl-1075, 1:1000); HuC/D (mouse, 1:20, Invitrogen, Grand Island, NY), which labels retinal ganglion cells and amacrine cells; Prox1 (rabbit, 1:2,000, Millipore, Billerica, MA), which recognizes horizontal cells; anti-PKCα (mouse, 1:100; cat. no. sc-17769, SantaCruz Biotechnology), which labels bipolar cells; glutamine synthetase (mouse, 1:500, BD Biosciences, Franklin Lakes, NJ) which labels Müller glia; anti-Blue and anti-UV opsin antibodies (rabbit, 1:1,000), generously provided by D. Hyde (University of Notre Dame); and 1D4, which labels mouse rhodopsin (Santa Cruz, sc-5743). Alexa fluor conjugated secondary antibodies (Invitrogen, Grand Island, NY) and Cy-conjugated secondary antibodies (Jackson ImmunoResearch, West Grove, PA) were used at 1:250 dilution. In addition, for anti-mouse Chd7, anti-mouse Chd7 primary antibody was followed by signal amplification with goat anti-rabbit IgG horseradish peroxidase (1: 500) (Perkin Elmer Inc: NEF812001EA, Waltham, MA). The tyramide signal amplification plus Cy3 Kit (1:1500) (Perkin Elmer Inc: NEL744001KT, Waltham, MA) was then used for detection. Terminal deoxynucleotide transferase (TdT)-mediated dUTP nick end labeling (TUNEL) was performed on frozen retinal cryosections using the ApopTag Fluorescein Direct In Situ Apoptosis Detection Kit (Millipore, Billerica, MA). TUNEL staining was performed according to the manufacturer’s protocol. For imaging, sections were counterstained with 4’, 6-diamidino-2-phenylindole (DAPI, 1:10,000 dilution, Sigma-Aldrich). Images were obtained on a Nikon Eclipse Ti-U inverted fluorescent microscope (Nikon Instruments, Melville, NY) or a Leica SP8 DLS confocal microscope (Leica Microsystems, Buffalo Grove, IL). For all experiments, at least 10 zebrafish embryos and 3 mouse embryos/pups were analyzed per timepoint, and 3 separate biological replicates were performed for each experiment.

### 2.7 Data Analysis and Figure Construction

All counts and measurements were performed on retinal cryosections containing an optic nerve as a landmark. Zebrafish retina size and mouse retinal layers were measured using ImageJ software (https://imagej.nih.gov/ij/). For zebrafish photoreceptors, counts were obtained from three separate regions 100 μM wide: a dorsal region 50 μM from the retinal margin, a central region 50μM dorsal to the optic nerve, and a ventral region 50 μM from the retinal margin. For mouse cone photoreceptors, counts were obtained from two 500 μM-wide regions on either side of the optic nerve. These counts were totaled. Statistics were conducted using two-factor, unpaired t-test and one-way ANOVA using GraphPad software. P-values less than 0.05 were considered significant and are indicated by the following: *, p<0.05; **, p<0.01; and ***, p<0.001. Boxplots were generated using the ggplot2 package in R (version 3.6.2; https://www.R-project.org). All figures were constructed using Photoshop (Adobe version 22.0.0).

## 3. Results

### 3.1 Chd7 Expression in Developing Retina

#### 3.1.1 Chd7 is expressed in distinct subsets of differentiated retinal cell types

To date there have been no published data on the expression of Chd7 in the developing zebrafish retina. We investigated Chd7 expression using immunohistochemistry (IHC) with an anti-zebrafish Chd7 antibody on retinal sections taken at various timepoints during retinal development. At 24 and 48 hours post fertilization (hpf), when retinal progenitor cells are still actively proliferating and the first retinal ganglion and amacrine cells are becoming postmitotic, Chd7 was broadly expressed throughout the zebrafish retinal neuroepithelium (Fig. 1A-B). By 72 hpf, when the majority of retinal cell types have exited the cell cycle and differentiated, Chd7 expression remained strong in the ganglion cell layer (GCL) and in cells in the basal portion of the inner nuclear layer (INL), which are likely amacrine cells (Fig. 1C). Interestingly, strong expression of Chd7 was also observed throughout the outer nuclear layer (ONL) containing the cone and rod photoreceptor cell bodies. By 4 and 5 dpf, when the zebrafish larvae display active swimming and visual behaviors (such as searching for food and avoiding predators), Chd7 expression was reduced but still visible in the GCL and inner INL, and remained strong in the photoreceptors of the ONL. Chd7 was also continuously detected in the persistently neurogenic ciliary marginal zone (CMZ) at 3, 4, and 5 dpf (Fig. 1C-E). To investigate the ONL expression of Chd7 further, we performed IHC for Chd7 on retinal sections from transgenic reporter lines that fluorescently label cones and rods (TαC:GFP labels all cone photoreceptor subtypes; XOPs:GFP labels rod photoreceptors). Chd7 expression significantly overlapped with GFP-positive cone photoreceptors at 72 hpf and 5 dpf (Fig. 1F-G). In contrast, co-expression of Chd7 with GFP-positive rod photoreceptors was minimal at 72 hpf, but was clearly present in rods by 5 dpf (Fig. 1H-I). Taken together, these results demonstrate expression of Chd7 in retinal progenitor cells and a subset of fully differentiated retinal neurons, including the photoreceptors.

**Figure 1.**
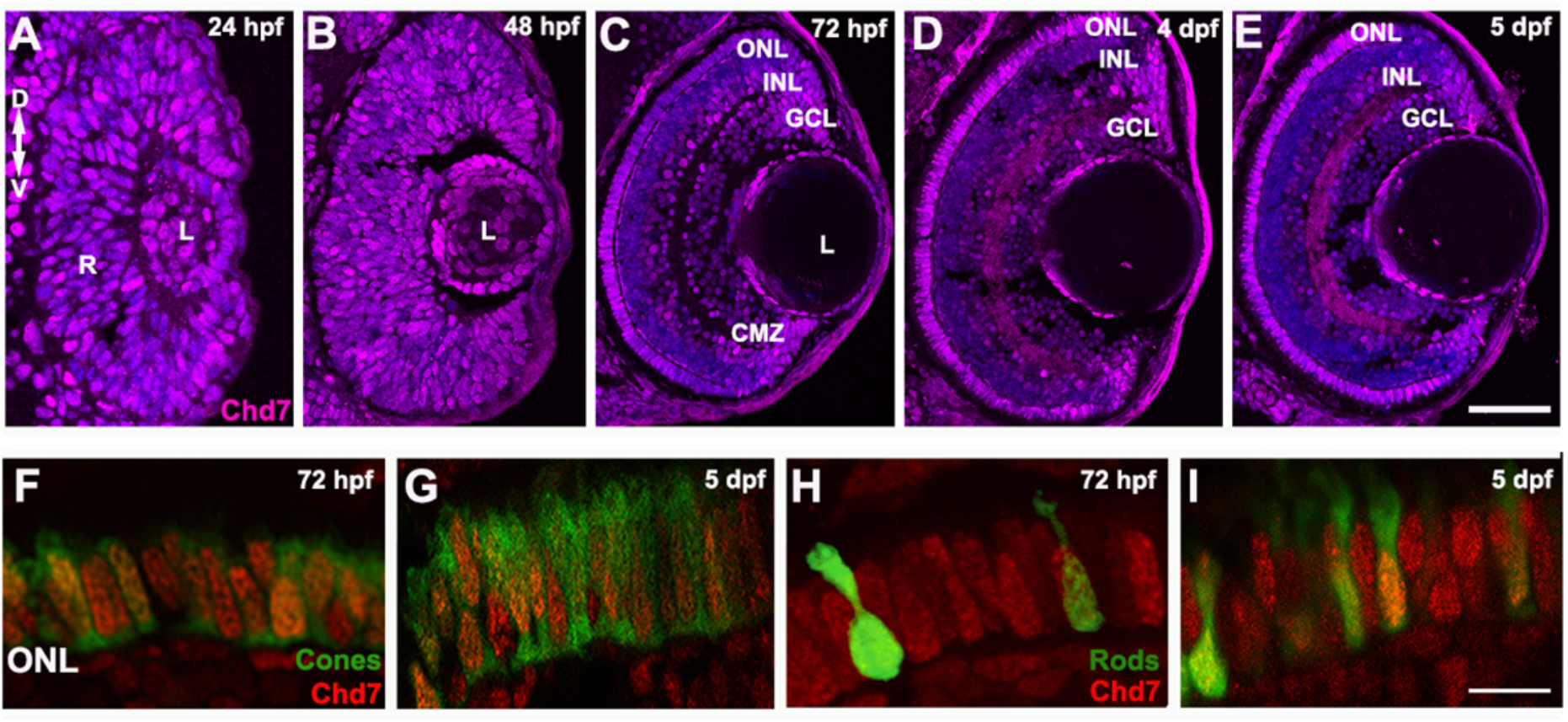
Chd7 is expressed in retinal progenitor cells and differentiated retinal cell types in zebrafish. (A-B) Broad expression of Chd7 is apparent in the developing retina at 24 and 48 hours post fertilization (hpf). (C) At 72 hpf, Chd7 is expressed throughout ganglion cell layer (GCL), some cells of inner nuclear layer (INL), sporadic cells of outer nuclear layer (ONL), and the ciliary marginal zone (CMZ). (D-E) Chd7 expression at 4 and 5 days post fertilization (dpf) shows reduction in GCL and INL expression and strong ONL expression. (F-G) At 72 hpf (F) and 5 dpf (G) Chd7 is co-expressed in GFP-positive cone photoreceptors marked by the TαC:GFP transgene. IHC for Chd7 in the XOPs:GFP transgenic line which fluorescently labels rod photoreceptors demonstrates that Chd7 has minimal co-expression with rod photoreceptors at 72 hpf (H) and increased co-expression at 5 dpf (I). ONL, outer nuclear layer; INL, Inner nuclear layer; GCL, ganglion cell layer; L, lens; CMZ, ciliary marginal zone; D, dorsal; V, ventral; R, retinal neuroepithelium; hpf, hours post fertilization; dpf, days post fertilization. Scale bars, 10μm in FI and 50μm in A-E.

#### 3.1.2 Mouse retinal CHD7 expression is similar to that of zebrafish

To determine whether CHD7 demonstrates similar expression patterns in the developing mouse retina, we performed IHC on mouse retinal sections at various embryonic and postnatal stages. CHD7 was expressed throughout the inner neuroblastic layer (INBL) in the developing mouse retina at embryonic day 15.5 when retinal progenitors are actively proliferating, and early cells of the retina are beginning to differentiate (Fig 2A). At E18.5 CHD7 expression was detectable in the newly formed ganglion cell layer (GCL), and CHD7 expression remained strong in the GCL at all subsequent timepoints (Fig. 2B-F). Sporadic expression of CHD7 was observed throughout the inner and outer neuroblastic layers (ONBL) at E18.5 and P1 (Fig. 2B-C). At P6, when retinal lamination was complete, CHD7 expression was prominent in the inner part of the INL but was not detectable in the ONL (Fig. 2D). This INL expression remained strong at P10 (when retinal progenitors are no longer present), and faint CHD7 expression also became evident in sporadic cells of the ONL (Fig. 2E). Interestingly, at P15 (just after eye opening) CHD7 expression remained strong in the GCL, numerous cells of the INL and was prominent in the distal portion of the ONL (Fig. 2F). The specificity of the CHD7 antibody signal was validated by observing expression in the *Chd7^Gt/+^* gene-trap mutant mouse line (Hurd et al., 2007), which demonstrated a markedly reduced signal at E15.5 and P15. The mouse expression data, along with the zebrafish expression pattern, suggests a conserved novel role for CHD7 in retinal progenitor cells and retinal neurogenesis and a previously undescribed expression and role in fully differentiated retinal cells.

**Figure 2.**
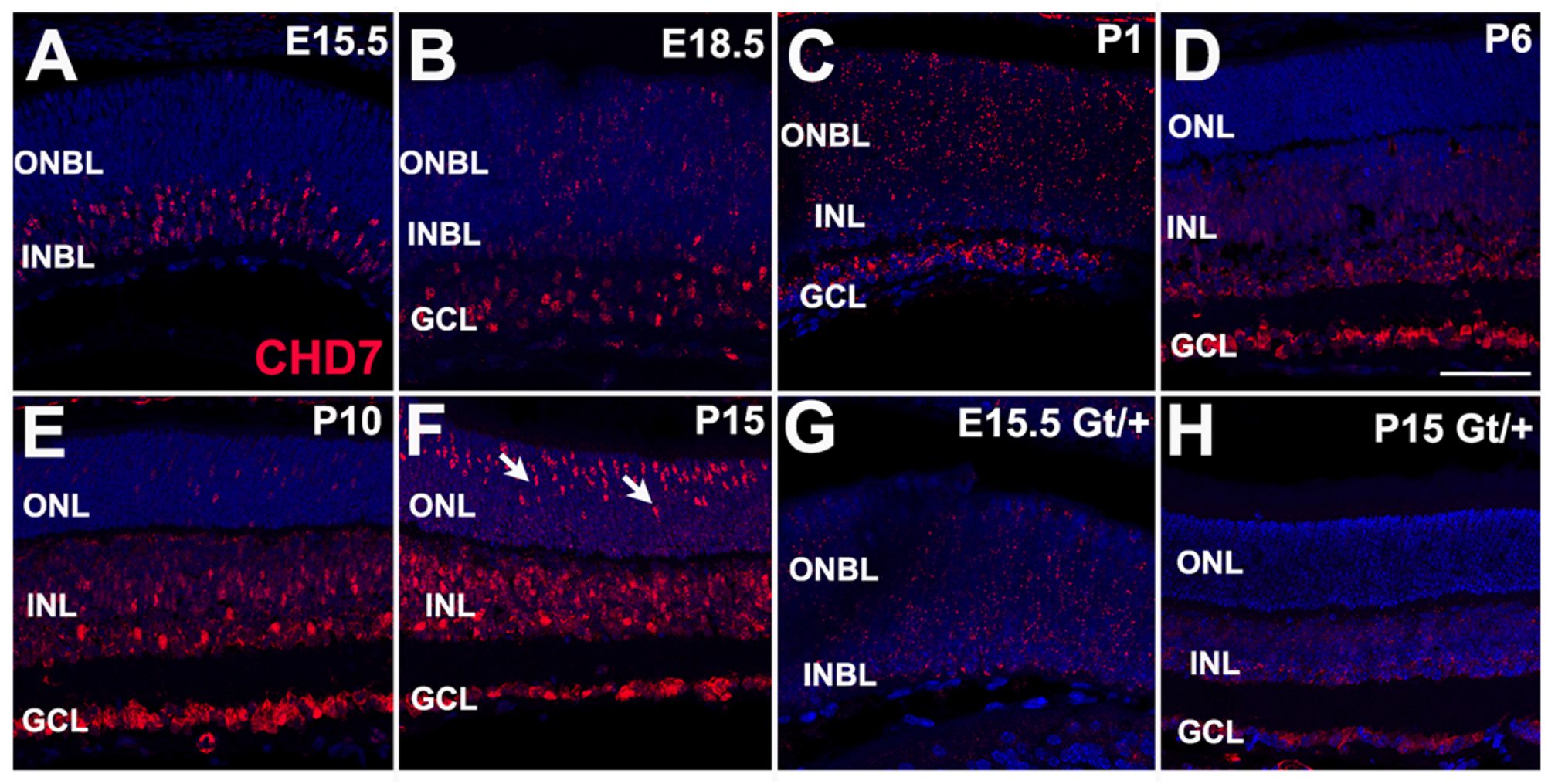
CHD7 is expressed in retinal progenitor cells and differentiated retinal cell types in mouse. (A) CHD7 expression is strong in the inner neuroblastic layer (INBL) at E15.5 (B) At E18.5, CHD7 expression continues in the INBL and is observed in sporadic cells of the outer neuroblastic layer (ONBL). (C-F) CHD7 expression at postnatal days P1, P6, P10, and P15. At P1 (C) expression is strong in the ganglion cell layer (GCL) and throughout the inner nuclear layer (INL) and ONBL. At P6 and P10 (D-E), CHD7 expression becomes restricted to the GCL and inner cells of the INL, and expression begins in a few cells of outer nuclear layer (ONL). By P15 (F), CHD7 expression remains strong in GCL, cells of the INL, and in a band of cells in the distal ONL. (G-H) CHD7 expression is significantly decreased in *Chd7^Gt/+^* mutant embryos (E15.5 and mice (P15). Arrows indicate expression in presumptive photoreceptor cells of the ONL. ONL, outer nuclear layer; INL, Inner nuclear layer; GCL, ganglion cell layer; ONBL, outer neuroblastic layer; INBL, inner neuroblastic layer. Scale bar is 100 μm for all panels.

#### 3.1.3 *CHD7* is expressed in the developing and mature human retina

To determine whether *CHD7* is expressed in the developing and adult human retina, we searched two publicly available data meta-analysis platforms. Using the eyeIntegration v1.05 platform which collates human eye tissue RNA-seq data with other human body tissues (Bryan et al., 2018), *CHD7* was shown to be globally expressed in fetal human retina and postnatal retina at levels equal to that of the cerebellum (Fig. S1A). Using the online PLatform for Analysis of scEiad (Single Cell Eye in a Disk) (Swamy et al., 2021), we generated an in-situ projection of *CHD7* expression in human adult retina, which indicates highest *CHD7* expression in cones, rods, and retinal ganglion cells with lower expression in Müller glia (Fig. S1B). In addition, when including data from fetal human retina and retinal organoids, expression of *CHD7* was detected in early and late retinal progenitor cells including photoreceptor precursors (Supplementary Table 2). These expression data, together with our zebrafish and mouse IHC results, strongly suggest a conserved role for CHD7 in retinal development across vertebrates.

### 3.2 Gross morphology of zebrafish *chd7* mutants

#### 3.2.1 Characterization of zebrafish *chd7* mutant alleles

To further explore the role of *chd7* in retinal development we obtained two different zebrafish mutant lines. The first line is a previously uncharacterized N-ethyl-N-nitrosourea (ENU) mutant generated by the Zebrafish Mutation Project at the Wellcome Sanger Institute (obtained from ZIRC – allele designation *sa19732*). This allele contains a point mutation in the helicase domain of Chd7 resulting in a premature stop codon and predicted truncated mutant protein of 1449 amino acids (compared to wildtype length of 3140 amino acids) (Fig. 3A). We confirmed loss of Chd7 protein expression in this line via western blot, which revealed a decrease in Chd7 expression in heterozygous animals and complete loss of Chd7 protein expression in homozygous mutant animals (Fig. S2A). The second mutant line is a previously characterized CRISPR mutant with a 2-bp deletion in exon 2 of *chd7*, resulting in a frameshift and early termination codon, and a predicted protein of only 43 amino acids (Cloney et al., 2018). CHARGE-relevant phenotypes previously described in this mutant line include pericardial edema, microphthalmia, abnormal curvature of the spine and cardiac and gastrointestinal defects (Cloney et al., 2018).

**Figure 3.**
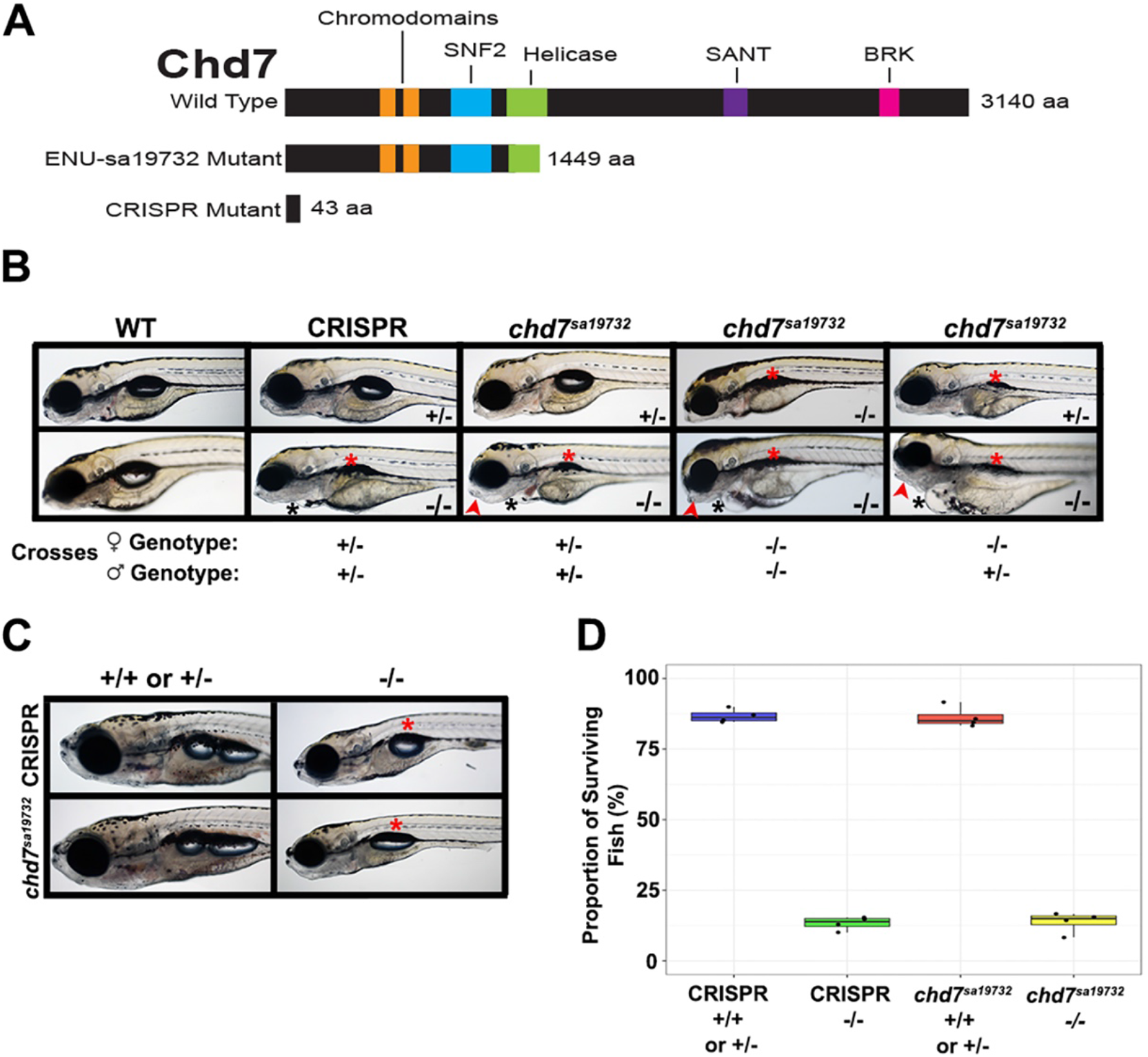
*chd7* mutant zebrafish display gross morphological defects and decreased survival. (A) Schematic of wildtype and predicted *chd7^sa19732^* and CRISPR mutant proteins with indicated protein domains. The *chd7^sa19732^* mutant protein is predicted to be 1449 aa and CRISPR mutant protein is predicted to be 43 aa compared to wild type length of 3140 aa. (B)*chd7* mutants display craniofacial, swim bladder, and pericardial abnormalities at 5 dpf compared to wildtype. (C) Gross morphology of WT and *chd7* mutants at 14 dpf; *chd7* mutants continue to display morphological defects and are smaller in size. Black asterisks indicate pericardial edema; red asterisks indicate lack of or smaller swim bladder. Red arrows indicate craniofacial abnormalities. (D) Proportion of surviving mutants at two months shows less than the expected 25% of homozygous mutants.

#### 3.2.2 Zebrafish *chd7* mutants display phenotypes observed in CHARGE syndrome

We first characterized the gross morphology of the *chd7^sa19732^* mutant compared to the previously described CRISPR mutant using light microscopy of progeny from heterozygous incrosses. At 5 dpf we observed a lack of swim bladder inflation, pericardial edema, microphthalmia, and craniofacial abnormalities in both the homozygous CRISPR mutant and the *chd7^sa19732^* mutant compared to wildtype and heterozygous larvae (Fig. 3B). At 14 dpf, both the homozygous *chd7^sa19732^* and CRISPR mutants remained significantly smaller than their heterozygous and wildtype siblings, and continued to display a smaller swim bladder, craniofacial, and spinal deformities (Fig. 3C). We also preformed intercrosses between the *chd7^sa19732^* and CRISPR mutants and observed similar phenotypes in compound heterozygotes (data not shown). We noticed a significant decrease in homozygous mutant survival, with only 13% of *chd7^-/-^* larvae surviving to adulthood (Fig. 3D). Nevertheless, the surviving adult homozygous mutants were fertile, and crosses of *chd7* mutant females and males produced clutches of *chd7* mutant offspring. The maternal zygotic *chd7* mutant progeny (from the *chd7^sa19732^* mutant line) displayed more severe pericardial edema, microphthalmia, and craniofacial abnormalities than the zygotic mutants (Fig. 3B). Furthermore, when homozygous mutant females were crossed with heterozygous males, heterozygous progeny displayed phenotypes similar to their homozygous mutant siblings, suggesting that the presence of maternal *chd7* mitigates the phenotypes of zygotic mutants. Due to the low numbers of homozygous *chd7* mutants that survive to adulthood, we characterized *chd7* zygotic mutants for the majority of this study.

#### 3.2.3 Loss of Chd7 results in microphthalmia

We next investigated overall retinal structure of *chd7* mutants at 5 dpf. Using retinal cryosections immunolabeled with the anti-Chd7 antibody along with nuclear staining, we observed a decrease of Chd7 expression in the retina of heterozygous *chd7* mutants, and confirmed complete loss of Chd7 expression in homozygous mutants at 5 dpf (Fig. 4A-F). We observed no significant defects in laminar organization nor evidence of retinal coloboma in either *chd7* mutant line at this stage. However, both CRISPR and *chd7^sa19732^* mutants were microphthalmic; we quantified relative eye size by measuring the retinal perimeter and normalizing that to the nose-to-otolith length to account for any gross larval size differences across genotypes. In both the CRISPR and *chd7^sa19732^* mutants, there was a significant difference between the normalized wildtype, heterozygous, and homozygous eye sizes (*p*<.05); homozygous mutants displayed the most severe reductions in eye size (*p*<.01 in the *chd7^sa19732^* mutant and *p*<.001 in the CRISPR mutant), suggesting an additive effect of each mutant allele on this phenotype. (Fig. 4G). This change in retinal size was also observed in the heterozygous *Chd7* mice, where there was a significant difference in the width of both the inner nuclear layer and outer nuclear layer compared to wildtype (Fig. S3A). These results suggest that loss of Chd7 in zebrafish results in microphthalmia but not coloboma (in zebrafish), recapitulating some of the ocular malformation defects observed in CHARGE syndrome.

**Figure 4.**
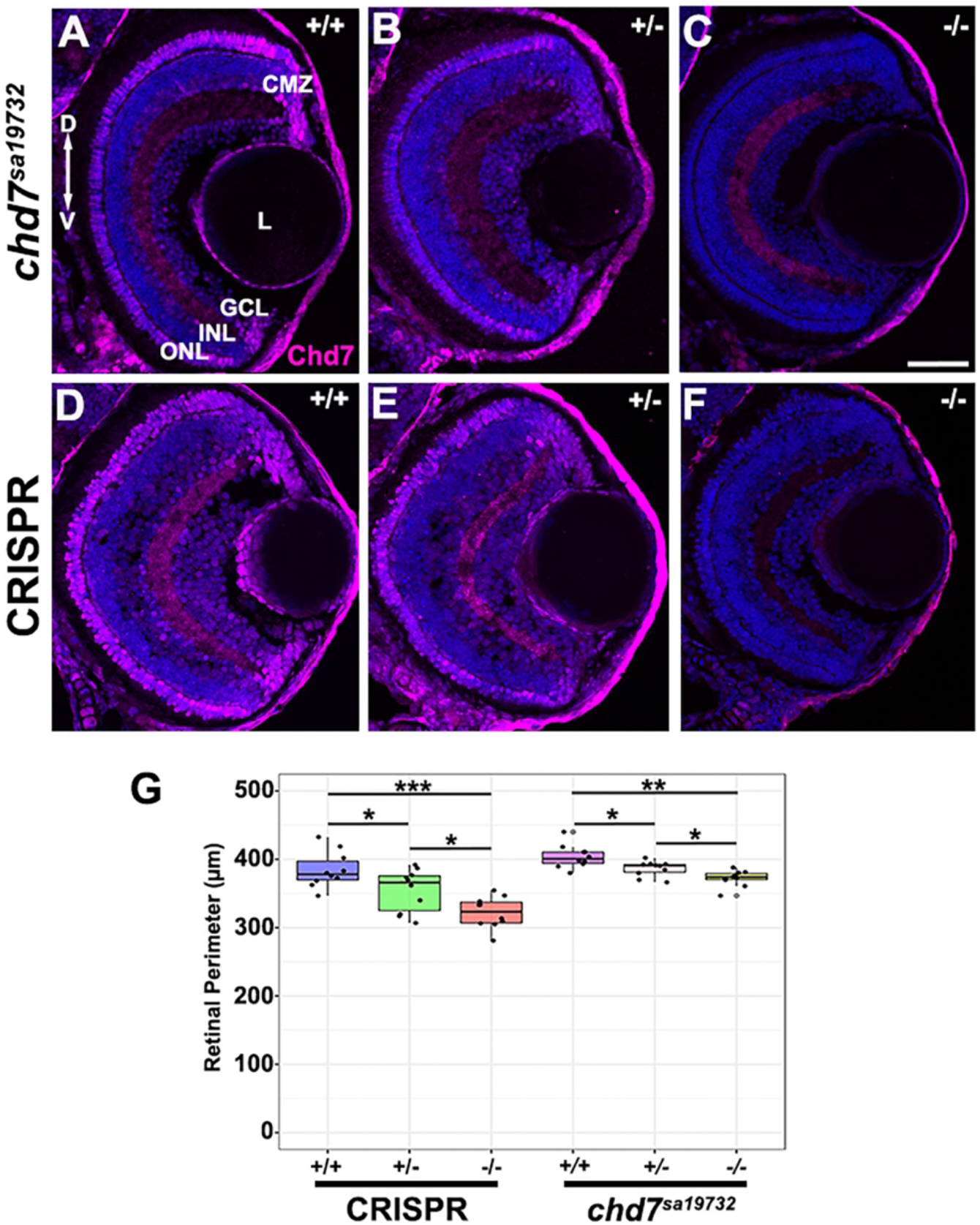
*chd7* mutants lack retinal Chd7 expression and display microphthalmia. (A-C). Chd7 expression in wild type, *chd7* heterozygous, and *chd7* homozygous mutant retinas of *chd7^sa19732^* mutant line (D-F). Chd7 expression in wild type, *chd7* heterozygous, and *chd7* homozygous mutant retinas of CRISPR mutant line. Both lines showed decreased Chd7 expression in heterozygotes and a lack of Chd7 expression in homozygotes. (G) *Chd7* mutants have microphthalmia compared to wildtype zebrafish. Measurements taken from perimeter of retina normalized to nose-to-otolith length. (\**p* < 0.05, \*\**p*< 0.01, \*\*\**p*< 0.001). ONL, outer nuclear layer; INL, Inner nuclear layer; GCL, ganglion cell layer; L, lens; D, dorsal; V, ventral. Scale bar 50 μm.

### 3.3 Specific retinal cell defects in *chd7* zebrafish mutants

#### 3.3.1 *chd7* mutants display a decrease in cone but not rod photoreceptors

Because we observed strong expression of Chd7 in newly differentiated photoreceptors, we next wanted to determine whether loss of Chd7 resulted in any photoreceptor phenotypes. We quantified the number of rod and cone photoreceptors in wildtype and *chd7* CRISPR mutants at 5 dpf by IHC, using the Zpr1 antibody (which labels red-green double cones) and 4C12 (which detects rods) (Morris et al., 2005). In wildtype retinas, we observed a compact and continuous layer of red-green cones in the ONL at 5 dpf (Fig. 5A-A’). However, in *chd7* heterozygous and homozygous mutant retinas we observed gaps in double cone spacing across the entire ONL (Fig. 5B-C, Fig. 5B’-C’). We quantified red-green double cone density by counting the number of Zpr1-labeled cells within three 100 μm regions: the dorsal retina (50 μm from the dorsal CMZ), central retina (50 μm dorsal to the optic nerve), and ventral retina (50 μm from ventral CMZ). This quantification confirmed that red-green cones are significantly reduced in both heterozygous and homozygous *chd7* mutants compared to wildtype retinas (52+/-4 in wt vs. 41+/-4 in *chd7* heterozygotes and 33+/-3 in *chd7* homozygous mutants; *p*<.001 for both comparisons; n=10 per genotype; Fig. 5G). Interestingly, we did not observe a similar reduction in rod photoreceptors in *chd7* mutants at this stage (Fig. 5D-F’, H). We confirmed that the loss of cones was not unique to the CRISPR *chd7* mutant by conducting the same IHC experiments in the *chd7^sa19732^* mutant. These mutants had a similar gap in red-green cones in the ONL compared to their wildtype siblings (Fig. S4A-C, A’-C’), although the reduction was smaller than that observed in the CRISPR mutant (Fig. S4G). As with the CRISPR mutants, rod photoreceptor number in the *chd7^sa19732^* mutant was not significantly reduced compared to wild type retinas (Fig. S4D-F, D’-F’, and Fig. S4H). Taken together, these results demonstrate that the loss of Chd7 causes a significant decrease in cone, but not rod, photoreceptor number.

**Figure 5.**
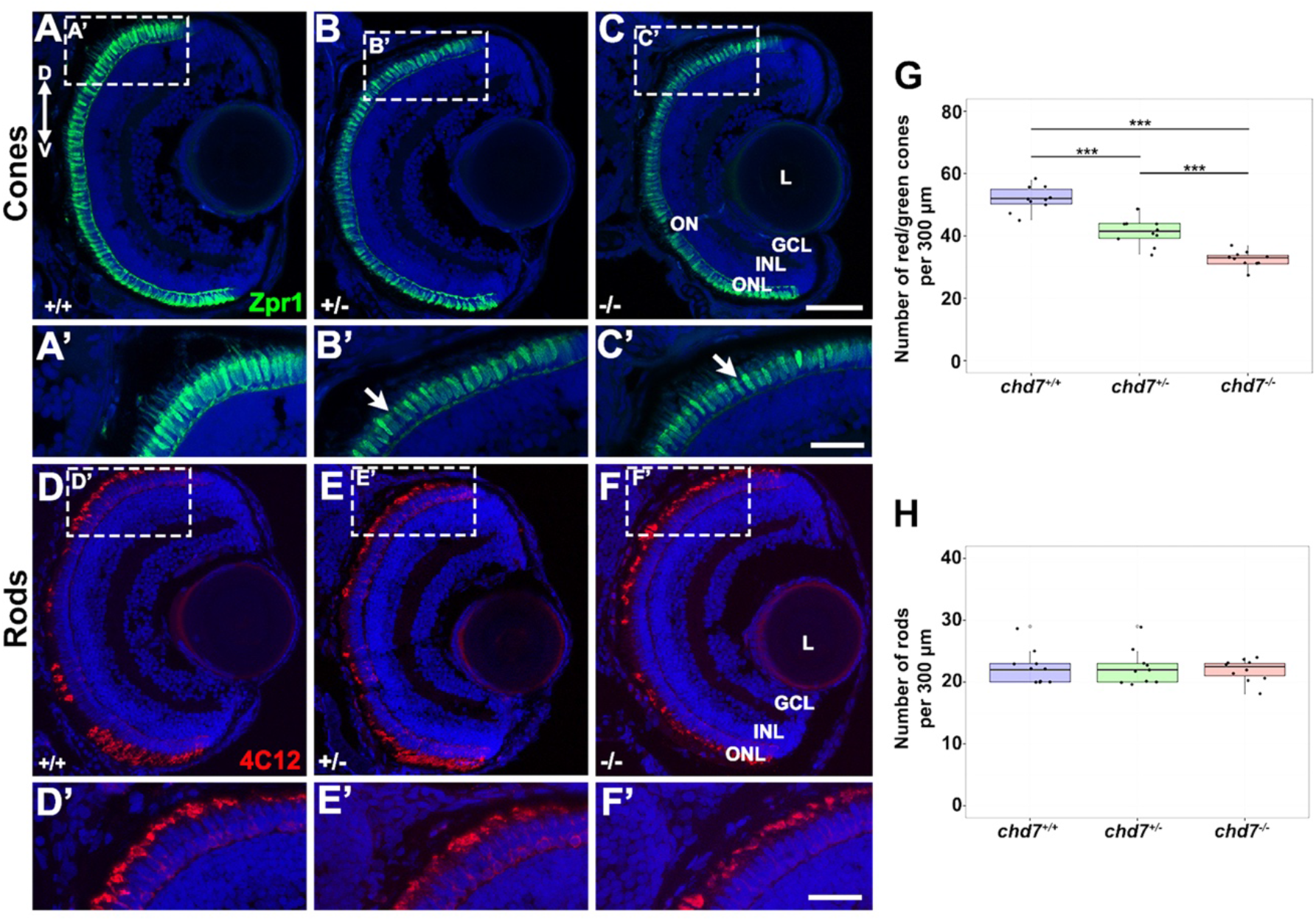
*chd7* mutant zebrafish have fewer cone photoreceptors. (A-C) Immunohistochemistry with a red-green cone antibody (Zpr1) in *chd7^+/+^* (A,A’), *chd7^+/-^* (B,B’), and *chd7^-/-^* (C,C’) retinal sections. Arrows indicate missing red-green cone photoreceptors. (D-F) Immunohistochemistry with a rod antibody (4C12) in *chd7^+/+^* (D,D’), *chd7^+/-^* (E,E’), and *chd7^-/-^* (F,F’) retinal sections. (G) Quantification confirms a decrease in red/green cones in *chd7^+/-^* and *chd7^-/-^* larvae compared to *chd7^+/+^*. (Number of cones per 300 μm; ANOVA followed by t-test; \*\*\**p*<.001). (H) No significant difference in rod photoreceptors between in *chd7^+/+^, chd7^+/-^*,and *chd7^-/-^* larvae (Number of rods per 300 μm). ONL, outer nuclear layer; INL, Inner nuclear layer; GCL, ganglion cell layer; L, lens; ON, optic nerve; D, dorsal; V, ventral. Scale bars 25 μm in A’-F’ and 50 μm in A-F.

#### 3.3.2 Cone and rod outer segments are truncated in *chd7* mutants

In addition to the reduced numbers of cones, we noticed a decrease in antibody labeling of the photoreceptor outer segments in *chd7* mutants. To further investigate this, we specifically labeled rod and cone outer segments using the 1D1 antibody for rod outer segments (Fadool, 2003) and peanut agglutinin (PNA) for cones (Blanks and Johnson, 1984). Whereas in 5 dpf wildtype retinas, we observed elongated rod outer segments that intercalated with the retinal pigment epithelium (RPE), in both the heterozygous and homozygous *chd7* mutants rod outer segment length appeared severely reduced (Fig. 6A-C). We quantified this reduction by averaging the length of 1D1-positive outer segments from 10 rods per eye in a minimum of 30 larvae of each genotype; this analysis confirmed a significant difference between wildtype and mutant rod outer segment length (14.4+/-0.86 μm in wt vs. vs. 6.6+/-0.47 μm in *chd7* heterozygotes and 5.6+/-0.25 μm in *chd7* null mutants; *p*<.001 for both comparisons; n=30 for each genotype; Fig. 6G). Similarly, we observed a significant decrease in the length of PNA-positive cone outer segments of *chd7* heterozygotes and homozygous mutants, with the truncation appearing most severe in *chd7*^-/-^ retinas (Fig. 6D-F). Quantifying the length of PNA-positive cone outer segments across genotypes revealed a reduction in outer segment length of 45% in heterozygous and 66% in homozygous *chd7* mutant cones compared to wild type (7.6+/- 0.31 μm in wt vs. 4.1+/-2.5 μm in *chd7* heterozygotes and 2.6+/-0.26 μm in *chd7* mutants; *p*<.001 for both comparisons; n=30 for each genotype; Fig. 6H). To determine whether all zebrafish cone subtypes were affected by the loss of Chd7, we performed immunohistochemistry with UV and blue cone opsin-specific antibodies. We observed similar reductions in blue and UV cone outer segments in *chd7^+/-^* and *chd7^-/-^* retinas compared with wild type retinas (Fig S5A-F). To evaluate whether the photoreceptor phenotypes persisted beyond the larval stage of development, we observed retinas of wildtype and *chd7* mutant larvae at 14 dpf. We detected similar photoreceptor phenotypes in *chd7* mutant retinas at this stage, including fewer red-green cones and truncated cone and rod outer segments (Fig. S6A-J). Taken together, these results demonstrate that that loss of Chd7 disrupts outer segment morphogenesis or maintenance in all zebrafish photoreceptors, and these defects persist into the juvenile stage.

**Figure 6.**
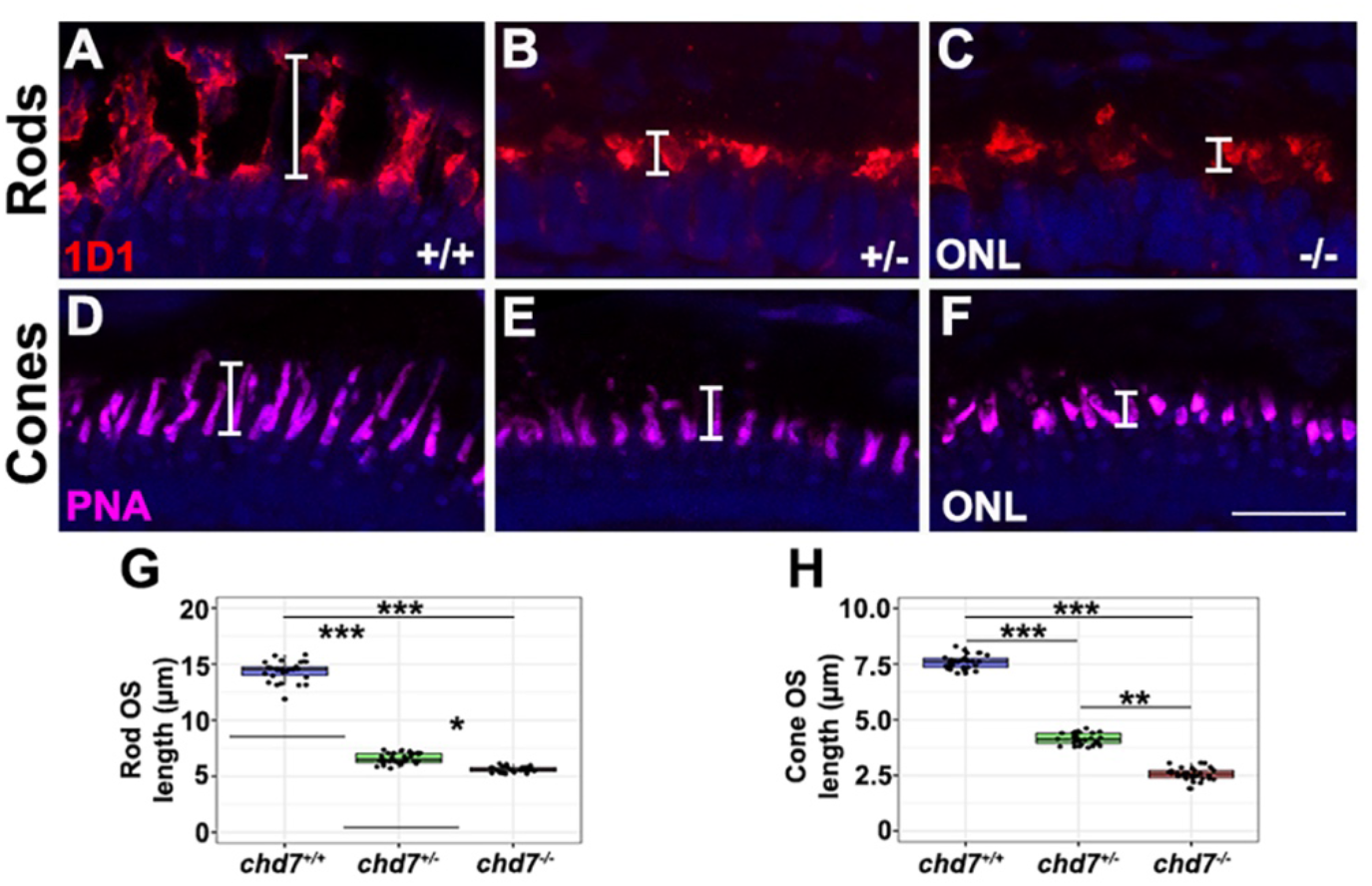
*chd7* mutant zebrafish display truncated photoreceptor outer segments. (A-C) Immunohistochemistry with a rhodopsin antibody (1D1) in *chd7^+/+^* (A), *chd7^+/-^* (B), and *chd7^-/-^* (C) retinal sections. (D-E) PNA labeling of cone outer segments in *chd7*^+/+^(D), *chd7*^+/-^ (E), and *chd7^-/-^* (F) retinal sections. (G) Average rod outer segment length in *chd7^+/-^* and *chd7^-/-^* larvae is significantly decreased compared to *chd7^+/+^*. (H) Cone outer segments also display a significant decrease in length in *chd7^+/-^* and *chd7^-/-^* larvae compared to *chd7^+/+^* larvae. Each data point represents the average measurement for an individual eye, with a minimum of 10 cells measured per eye. Brackets represent region of length measurement for individual outer segments. (\**p* < 0.05, \*\**p*< 0.01, \*\*\**p*< 0.001). ONL, outer nuclear layer. Scale bar, 15 μm.

#### 3.3.3 Other retinal cell types are unaffected in *chd7* CRISPR mutants at 5 dpf

To determine if the loss of Chd7 affects other retinal cell types we performed immunohistochemistry with cell-type specific antibodies for ganglion cells, amacrine cells, bipolar cells, horizontal cells, and Müller glia at 5 dpf. Using HuC/D (ganglion and amacrine cells), PKCa (bipolar cells), Prox1 (horizontal cells), and glutamine synthetase (Müller glia), we observed that all cell types were present and there was no obvious decrease in number or morphology of cells in *chd7* mutant retinas compared to wildtype retinas (Fig. S7A-L). While more subtle differences may be identified with further examination, these results suggest that the loss of Chd7 has the strongest impact on the number and morphology of photoreceptors.

#### 3.3.4 Apoptosis is not increased in *chd7* mutant retinas at 5 dpf

To determine whether the reduced numbers of cone photoreceptors in *chd7* mutants are the result of a decrease in their survival, we performed TUNEL labeling on retinal cryosections at 5 dpf in wildtype and *chd7* CRISPR zebrafish mutants to identify apoptotic cells. We observed no significant difference in number of TUNEL+ cells between retinal sections of wildtype and mutant larvae (Fig. S8A-C). While we cannot exclude the possibility that loss of Chd7 causes apoptosis in the retina at earlier developmental timepoints, this result suggests that cone photoreceptors of *chd7* mutants are not reduced via cell death at this stage.

#### 3.3.5 Maternal zygotic *chd7* mutants display more severe photoreceptor defects

Given the more severe gross morphological defects detected in maternal zygotic (MZ) *chd7* zebrafish mutants, we were interested in determining whether the eye size and photoreceptor defects were also more severe in *chd7* MZ mutants. Using the same morphometric and immunohistochemical methods as described above, we detected a significant decrease in eye size in *chd7* MZ mutants compared to zygotic mutants (Fig. S9E). We also found that MZ mutants have fewer red-green cones than zygotic mutants (Fig. S9A-A’), but the number of rod photoreceptors was again not significantly different (Fig. S9C-D, Fig. S9G). Taken together, this indicates that maternal Chd7 not only contributes to ocular morphogenesis and growth but is also important for photoreceptor development.

### 3.4 *Chd7^Gt/+^* mouse mutants display a loss of cone photoceptors and truncated rod and cone outer segments

Finally, to investigate whether CHD7 is required for proper photoreceptor development in a mammalian model, we examined photoreceptor number and outer segment morphology in *Chd7^Gt/+^* retinas. We used the 1D4 antibody to detect rhodopsin and PNA to label cone photoreceptor outer segments. At P6, when retinogenesis is complete in the central retina and outer segments are beginning to mature, we observed nascent rod and cone outer segments in both wildtype and *Chd7^Gt/+^* mouse retinas, although there were fewer PNA-positive cells in the *Chd7^Gt/+^* retina (Fig. 7A-D). By P15, we detected shorter rod outer segments in the *Chd7^Gt/+^* retina, and no discernable difference in rod density (Fig. 7E-F). For the cone photoreceptors, we detected a significant (40%) reduction in the number of cones in *Chd7^Gt/+^* retinas, based on total counts obtained from two 500 μm regions on either side of the optic nerve and excluding regions with evident coloboma (83 +/-2 cones in wt vs. 50+/-5 in *Chd7*^Gt/+;^ n=3 per genotype; p<.001; Fig. 7G-I). *Chd7^Gt/+^* cone outer segment length was also reduced relative to wild type cones. Overall, these results suggest that Chd7 has a conserved role in photoreceptor development and outer segment morphogenesis.

**Figure 7.**
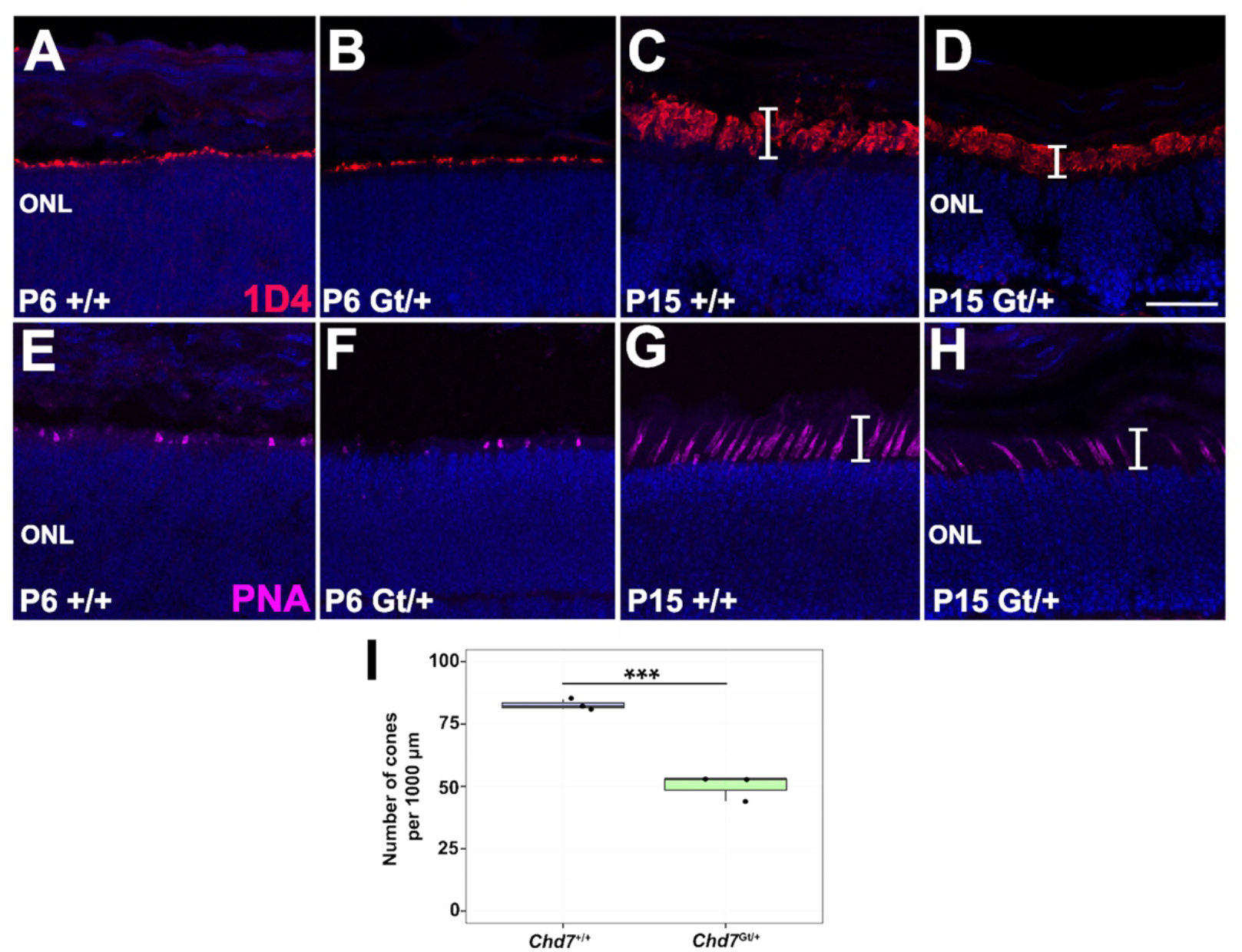
*Chd7* heterozygous mutant mice have fewer cones and truncated outer segments. (A-D) Immunohistochemistry with rhodopsin antibody (1D4) in *Chd7^+/+^* and *Chd7^Gt/+^* P6 and P15 retinal sections. (E-H) PNA labeling in *Chd7^+/+^* and *Chd7^Gt/+^* P6 and P15 retinal sections. Brackets indicate region of outer segments, which are shorter in *Chd7^Gt/+^* retinas. (I) Quantification of cone cell number shows a significant decrease in *Chd7^Gt/+^* compared to *Chd7^+/+^* at P15 (number of cones per 1000 uM, \*\*\**p* <.001). ONL, outer nuclear layer. Scale bar, 25 μm.

## 4. Discussion

Most studies on the ocular complications of CHARGE syndrome have focused on the early ocular morphogenesis defects – microphthalmia, coloboma, and optic nerve anomalies – that are associated with this disorder (Bajpai et al., 2010; Gage et al., 2015). Given that CHD7 has a demonstrated, critical role in neurogenesis in the CNS as well as in auditory and olfactory sensory neurons (Feng et al., 2017; Hurd et al., 2010; Layman et al., 2011, 2009), it is important to investigate whether CHD7 also contributes to the development of retinal neurons.

In this study, we demonstrated that Chd7 is expressed in zebrafish and mouse retinal progenitor cells, as well as in the persistently neurogenic ciliary marginal zone of the zebrafish retina. These expression data are in line with previously reported roles of CHD7 in neurogenesis. CHD7 and its chromatin remodeling function have been implicated in both the central and peripheral nervous systems. In the central nervous system, CHD7 has been shown to promote development of neurons in the brain, cortex, and cerebellum and in the neural stem cells of the subventricular zone (SVZ) of the lateral ventricle (LV) and the subgranular zone (SGZ) of the dentate gyrus (DG) in the adult hippocampus (Feng et al., 2013; Whittaker et al., 2017). In the peripheral nervous system, disruption of CHD7 leads to defects in neuron development of the olfactory and auditory system (Hurd et al., 2010; Layman et al., 2009). Furthermore, loss of CHD7 in mouse embryonic stem cells results in defects in neuron differentiation including disrupted neuronal complexity and length as a consequence of disturbed expression of pro-neural genes (Yao et al., 2020). Our studies suggest CHD7 may have a similar role in the specification and differentiation of various cell types in the retina.

We also found that CHD7 is expressed in terminally differentiated retinal cells in zebrafish, mouse, and humans, including ganglion cells, amacrine cells, and photoreceptor cells. To our knowledge, this expression pattern in post-mitotic retinal neurons has not been previously described and suggests that CHD7 has important functions in these newly differentiated cell types that bears further investigation.

In zebrafish and mouse models of CHD7 deficiency, we observed previously reported phenotypes of craniofacial defects, heart defects, and microphthalmia – all phenotypes relevant to CHARGE syndrome (Balow et al., 2013; Cloney et al., 2018; Hurd et al., 2007; Jacobs-Mcdaniels and Albertson, 2011; Liu et al., 2018). Interestingly, although coloboma was evident in mouse *Chd7* heterozygotes, we did not observe coloboma in either of the zebrafish mutant lines. This lack of coloboma in the zebrafish mutants could be due to expression of wildtype maternally provided *chd7* mRNA in zygotic mutants; however, this seems unlikely since there were no colobomas in *chd7* maternal zygotic mutants. It is possible that the rapid window of ocular morphogenesis in zebrafish compared to mammals allows for optic fissure closure to occur without deleterious effects of Chd7 loss in the developing eye. However, zebrafish *chd7* mutants display microphthalmia, suggesting that loss of Chd7 does have negative consequences even at early stages of zebrafish eye development. Alternatively, there may be another Chd family member (such as Chd5, which is expressed in the eye) that can compensate for the loss of Chd7 in zebrafish with respect to optic fissure fusion (Bishop et al., 2015; El-Brolosy et al., 2019). It is also possible that the molecular mechanism of choroid fissure closure is different between species, with Chd7 being required for certain steps of this process in the mouse but not in zebrafish.

One of the most intriguing findings of our study is that in both zebrafish and mouse *Chd7* mutants, we observed previously unreported photoreceptor phenotypes. These phenotypes include a significant decrease in cone photoreceptors and truncation of outer segments in both rod and cone photoreceptors. It is important to note that these retinal phenotypes were present in heterozygous as well as homozygous *chd7* mutants. The CRISPR mutant line used in our study was also previously reported to display gastrointestinal motility defects in both heterozygous and homozygous mutants. Taken together, these results suggest that loss of Chd7 in zebrafish results in haploinsufficient phenotypes similar to human CHARGE syndrome (Cloney et al., 2018), and the eye phenotypes further establish zebrafish as a model to investigate the pathogenesis of CHARGE syndrome-associated ocular defects.

Outer segments are essential for the phototransduction process, and thus are vital for vision. Given the lack of elongation of rod and cone outer segments in *Chd7* mutants of both zebrafish and mice, we predict that CHD7 plays a role in the development and/or maintenance of the unique photoreceptor outer segment structure. Photoreceptor outer segments, also known as photoreceptor cilia, are similar to other non-motile cilia and require many of the same abundant proteins for structure and function (Liu et al., 2007). *Chd7^Gt/+^* mice have previously been reported to have inconsistent and patchy labeling of olfactory cilia compared to wildtype mice (Layman et al., 2009). Taken together, we hypothesize that CHD7 may be required for the expression of one or multiple cilia genes and perhaps those that are essential for length of cilia or outer segment disk morphogenesis. Recently, CHD7 was also suggested to play a protective role in the auditory neurons and hair cells of the ear, and loss of CHD7 leads to oxidative stress-induced degeneration and sensorineural hearing loss (Ahmed et al., 2021). Given that hair cells and photoreceptors are both highly metabolically demanding sensory neurons, CHD7 could also be playing a role in oxidative stress response in photoreceptors. In future work it will be important to determine whether loss of CHD7 increases oxidative stress in the retina, and whether the altered outer segment morphology in CHD7 mutant photoreceptors results in reduced visual responses.

Although the expression pattern of CHD7 in the retina and photoreceptor phenotypes observed with loss of CHD7 strongly suggest that CHD7 is important for photoreceptor differentiation, to our knowledge photoreceptor defects have not yet been identified in individuals with CHARGE syndrome. This could be a result of the multifaceted nature of the other ocular malformations observed with *CHD7* mutations. The visual impairments resulting from coloboma, in addition to the complex clinical presentation of CHARGE syndrome, makes it challenging to perform normal vision assessments, potentially masking any accompanying retinal dystrophy. Moreover, photoreceptor defects may take years to become evident, and long-term studies on visual acuity in adults with CHARGE syndrome have not been reported. A recent study of visual function in 14 children with CHARGE syndrome found that all had some degree of visual impairment, even in the absence of structural malformations such as coloboma (Onesimo et al., 2021). These results suggest that the visual system disruptions associated with CHARGE syndrome may indeed involve retinal cell type defects in addition to problems with ocular morphogenesis. Thus, more work and novel assessments may be necessary in this patient population to identify potential underlying retinal cell pathology.

In conclusion, we have identified and characterized a novel expression pattern and role for CHD7 in retinal development. Future studies will help to identify the molecular targets of CHD7 in the developing retina and in newly differentiated photoreceptors, which should help to clarify the role of Chd7 in outer segment morphogenesis and maintenance. Taken together, our results suggest an important avenue of future investigation on the pathogenesis of visual system defects in CHARGE syndrome.

## Acknowledgements

We thank Brandi Bolton, Evelyn Turnbaugh, and Lucas Vieira Francisco for exceptional zebrafish care. We also thank members of the Morris lab for valuable input and technical assistance. We thank Dr. David Hyde (University of Notre Dame) for generously providing the zebrafish opsin antibodies and Dr. Jason Berman for proving the *chd7* CRISPR mutant line. We also thank Dr. Rob Hufnagel (National Eye Institute) for guidance with the meta-analysis platforms.

---

**Figure S1.**
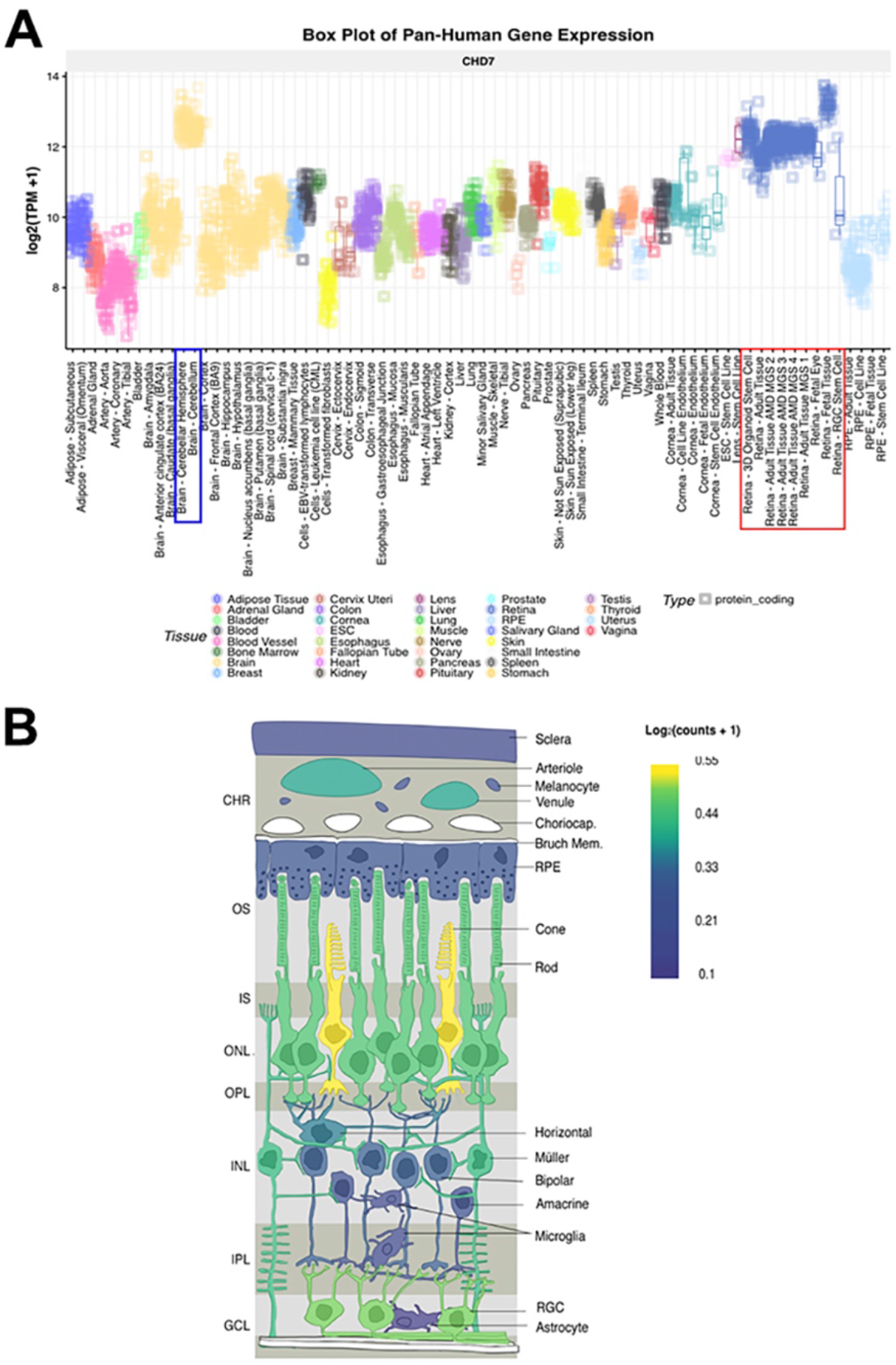
*CHD7* is expressed in cells of developing and adult human retina. (A) Box plot of pan-Human *CHD7* expression in fetal and adult retinal tissue (red box) compared to other tissues including cerebellum (blue box). Assembled with eyeIntegration v1.05 platform (Bryan et al., 2018). (B) In-situ projection from published human single cell RNAseq datasets. Assembled with PLatform for Analysis of scEiad (Swamy et al., 2021). Expression is highest in cones, rods, and retinal ganglion cells with lower expression in Müller glia.

**Figure S2.**
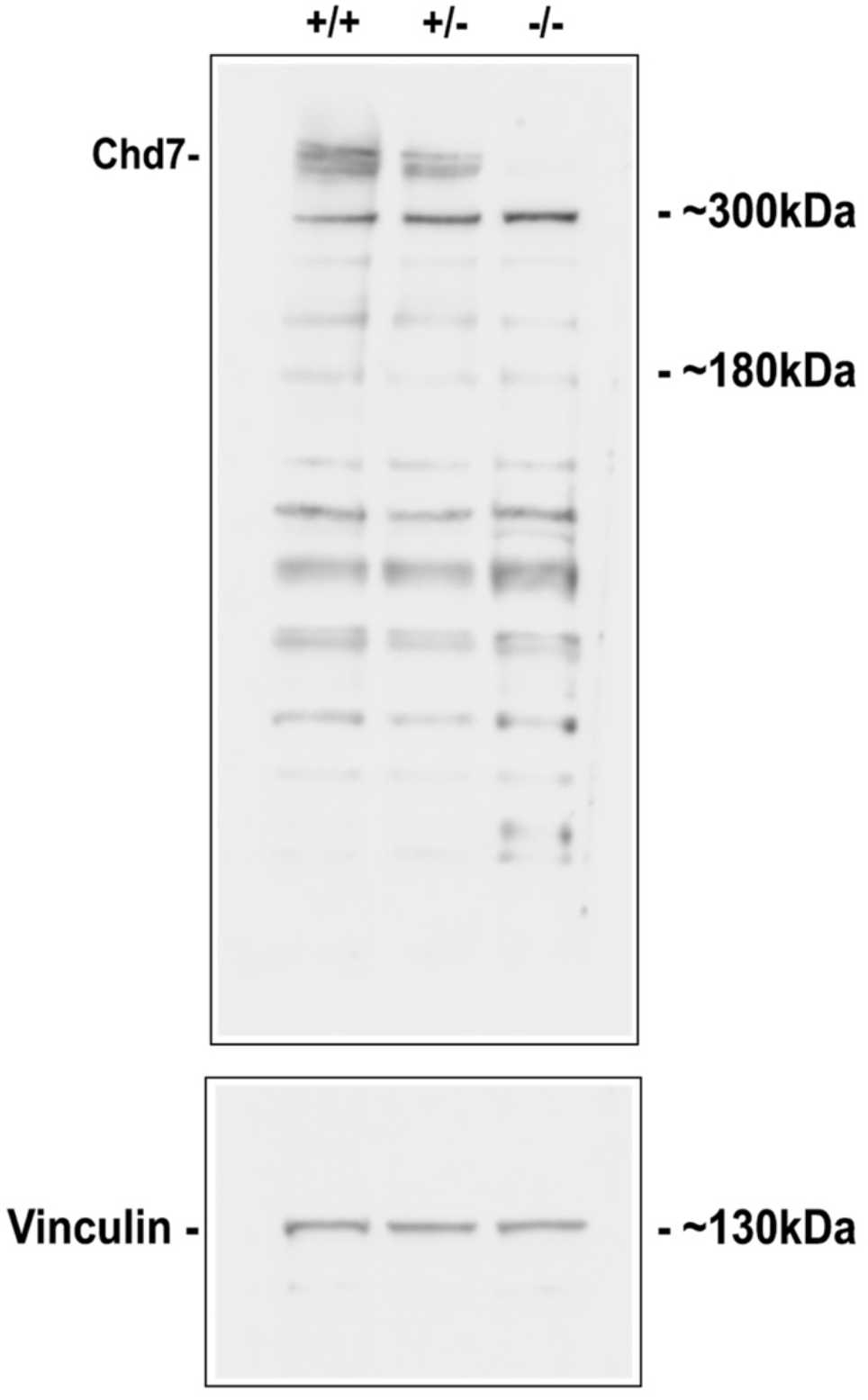
Loss of Chd7 protein expression in *chd7 chd7^sa19732^* mutant. Western blot of *chd7^+/+^, chd7^+/-^, chd7^-/-^* embryo lysate. Full-length Chd7 is 349 kDa. Lower blot is Vinculin loading control.

**Figure S3.**
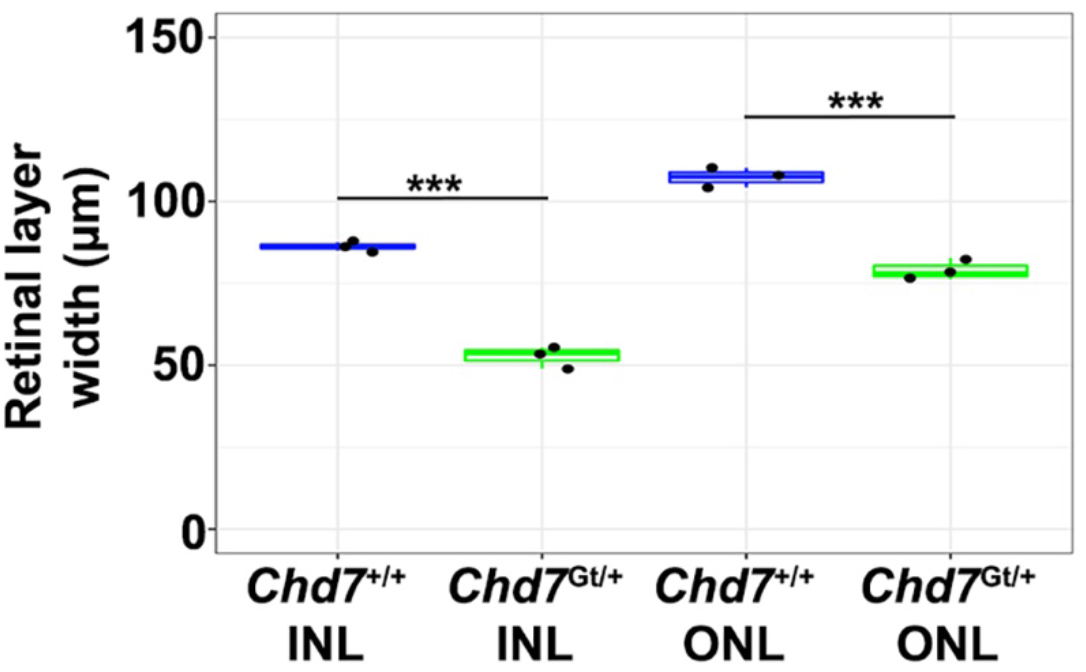
Retinal layers are smaller in *Chd7^Gt/+^* mutant mice compared to wild type. Width of inner and outer nuclear layer measurements between *Chd7^Gt/+^* mutant mice and *Chd7^+/+^* mice at P15 demonstrate significant decrease in *Chd7* heterozygous mutants. (t-test; n=3; \*\*\**p*< 0.001).

**Figure S4.**
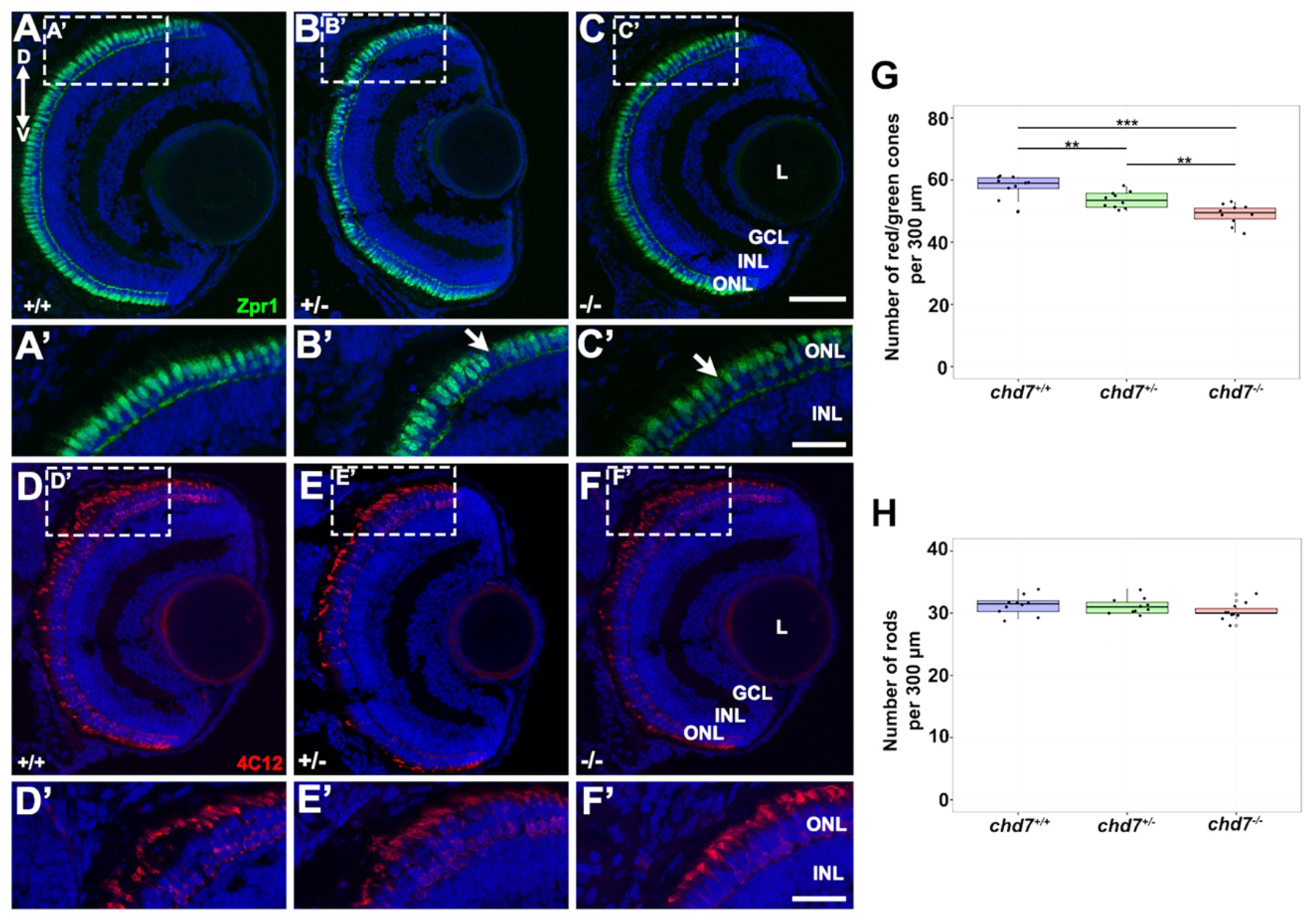
*chd7 chd7^sa19732^* mutant zebrafish have less cone photoreceptors. Immunohistochemistry with a red-green cone antibody (Zpr1) in *chd7^+/+^* (A,A’), *chd7^+/-^* (B,B’), and *chd7^-/-^* (C,C’) retinal sections. Arrows indicate missing red-green cone photoreceptors. (D-F) Immunohistochemistry with a rod antibody (4C12) in *chd7^+/+^* (D,D’), *chd7^+/-^* (E,E’), and *chd7^-/-^* (F,F’), retinal sections. (G) Quantification showed decrease in red/green cones in *chd7^+/-^* and *chd7^-/-^* larvae compared to *chd7^+/+^* (number of cones per 300 μm; ANOVA followed by t-test; n=10; \*\*\**p* <.001). (H) Quantification showed no difference in rod photoreceptors between in *chd7^+/+^, chd7^+/-^*, and *chd7^-/-^* larvae (number of rods per 300 μm). ONL, outer nuclear layer; INL, Inner nuclear layer; GCL, ganglion cell layer; L, lens; ON, optic nerve; D, dorsal; V, ventral. Scale bars 25 μm for A’-F’ and 50 μm for A-F.

**Figure S5.**
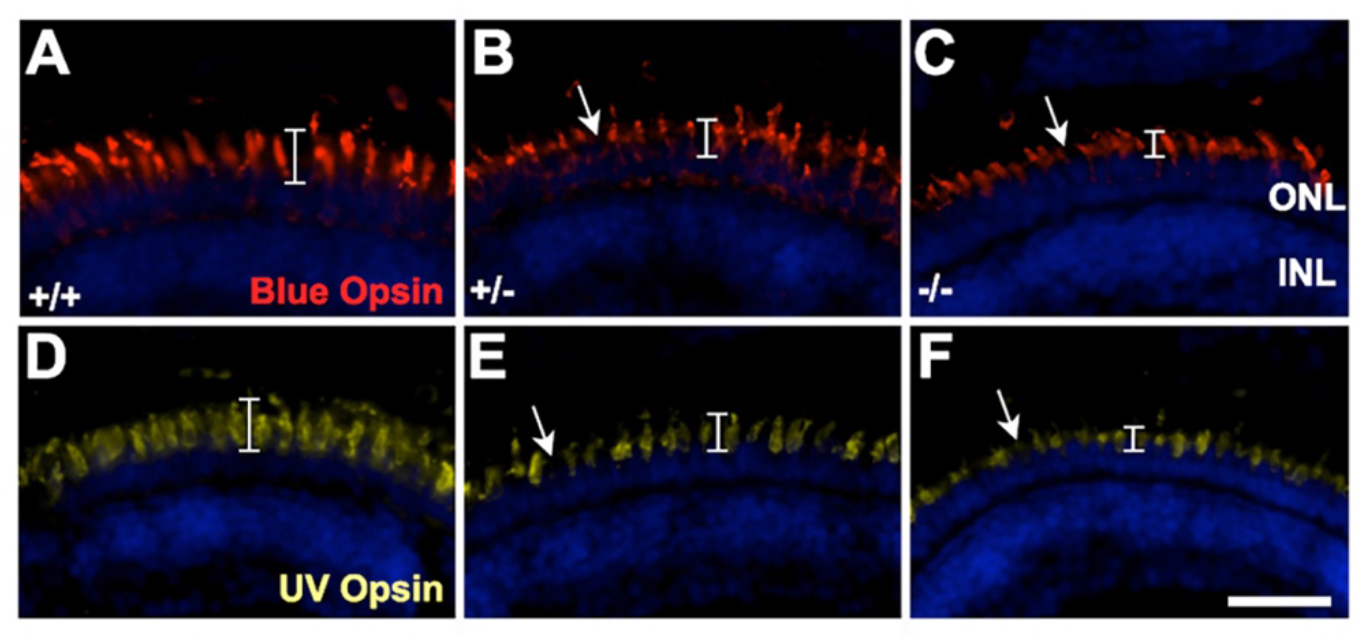
*chd7* CRISPR mutant zebrafish have reduced UV and blue cones. Immunohistochemistry using a blue (A–C), and UV (D–F) cone opsin antibody on *chd7^+/+^, chd7^+/-^*, and *chd7^-/-^* retinal cryosections shows a similar loss of UV and Blue cones as for red and green cones (Fig. 5). Arrows indicate missing cone photoreceptors. Brackets indicate length of outer segments. Scale bars, 25 μm.

**Figure S6.**
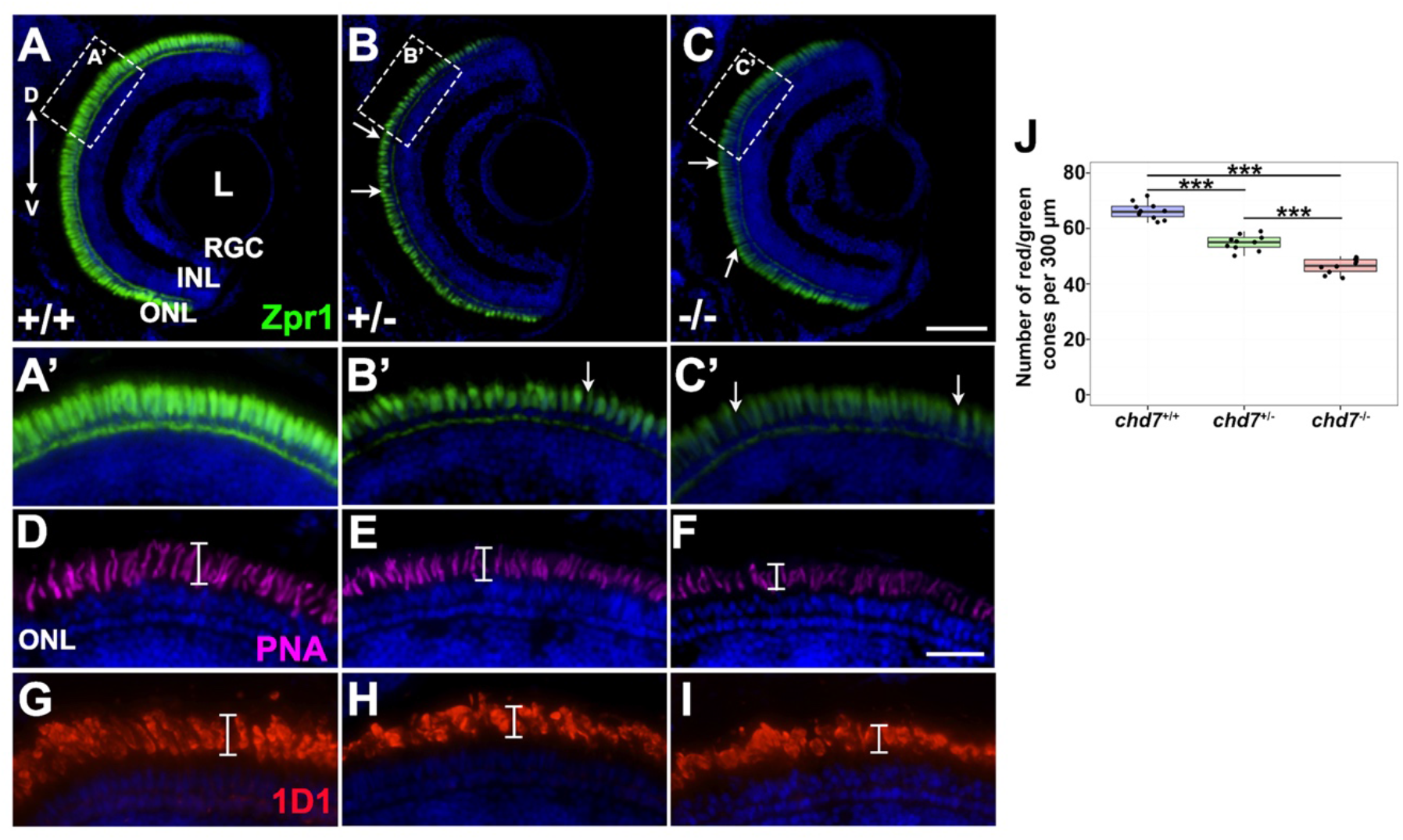
*chd7* CRISPR mutant zebrafish continue to display photoreceptor defects at 14 dpf. (A-C) Immunohistochemistry with a red-green cone antibody (Zpr1) in *chd7^+/+^* (A,A’), *chd7^+/-^* (B,B’), and *chd7^-/-^* (C,C’) retinal sections. Arrows indicate missing red-green cone photoreceptors. (D-F) PNA labeling for cones in *chd7*^+/+^(D), *chd7*^+/-^(E), and *chd7*^-/-^ (F) retinal sections. (G-I) Immunohistochemistry with a rhodopsin antibody (1D1) in *chd7^+/+^* (G), *chd7^+/-^* (H), and *chd7^-/-^* (I) retinal sections. (J) Quantification shows decrease in red/green cones in *chd7^+/-^* and *chd7^-/-^* larvae compared to *chd7^+/+^* (number of cones per 300 μm; ANOVA followed by t-test; n=10; \*\*\**p* <.001. ONL, outer nuclear layer; INL, Inner nuclear layer; GCL, ganglion cell layer; L, lens; D, dorsal; V, ventral. Scale bars 25 μm in A’-C’, D-I; 50 μm in A-C.

**Figure S7.**
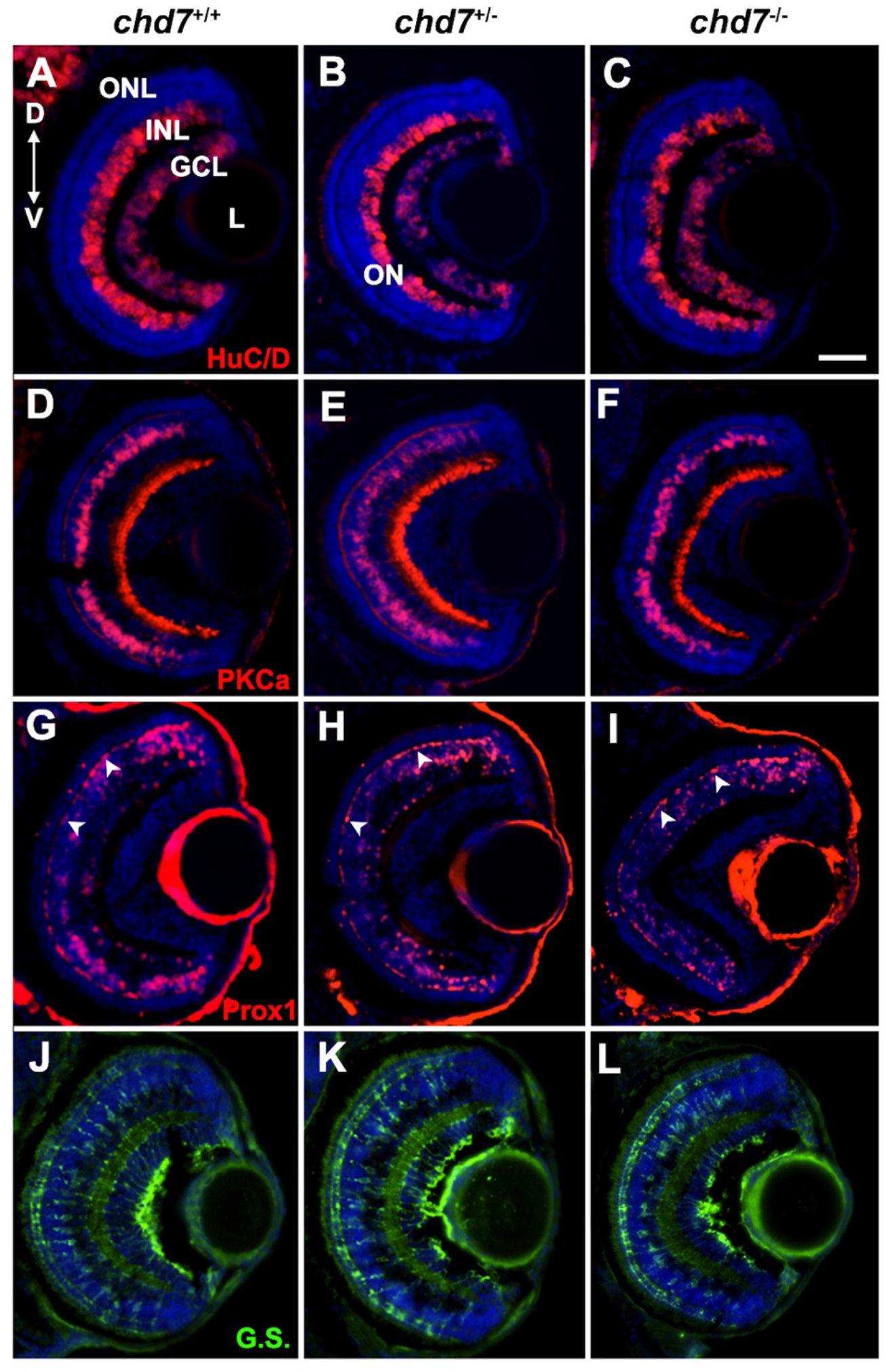
Other retinal cell types are unaffected in *chd7* CRISPR mutant zebrafish. Immunohistochemistry on retinal sections from each genotype for amacrine and ganglion cells with HuC/D antibody (A-C); bipolar cells with PKCa antibody (D-F); horizontal cells (arrowheads) with Prox1 antibody (G-I); and Müller glia with glutamine synthetase antibody (J-L). ONL, outer nuclear layer; INL, Inner nuclear layer; GCL, ganglion cell layer; L, lens; ON, optic nerve; D, dorsal; V, ventral. Scale bar 50 μm.

**Figure S8.**
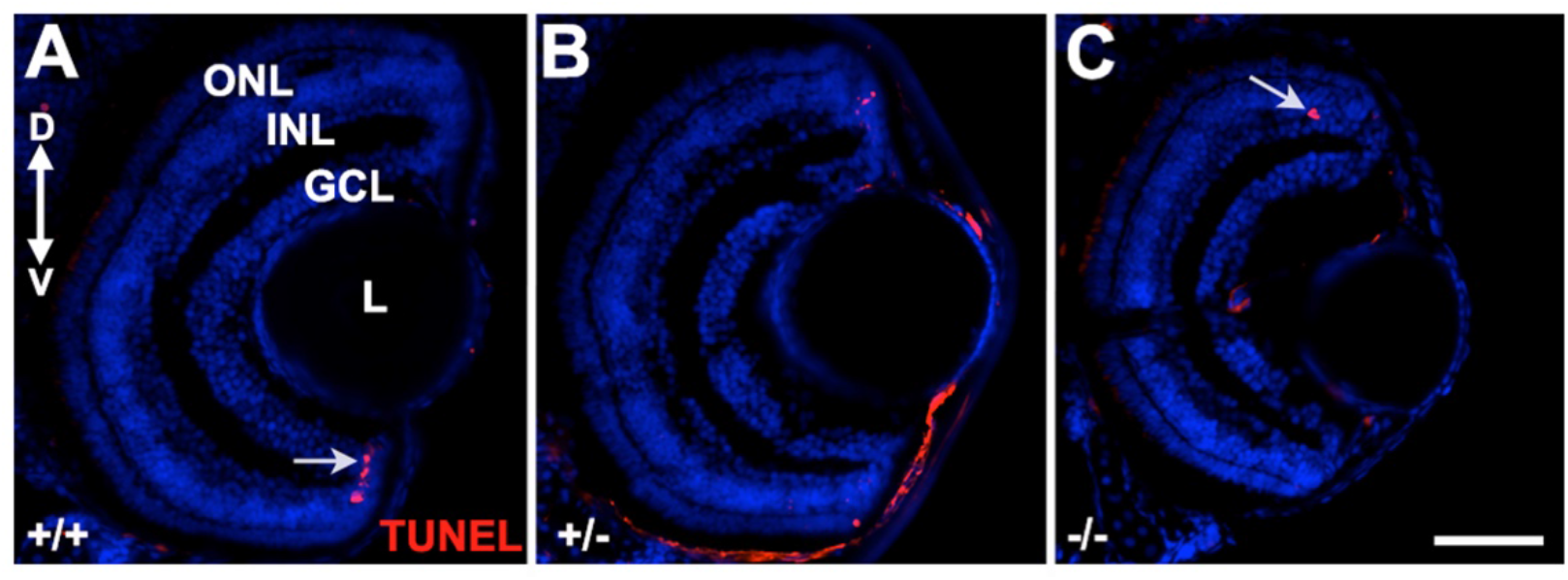
Apoptosis is not increased in *chd7* zebrafish mutant retinas at 5 dpf. (A–C) TUNEL staining in *chd7^+/+^, chd7^+/-^*, and *chd7^-/-^* retinal sections. Arrows indicate TUNEL+ cells. ONL, outer nuclear layer; INL, Inner nuclear layer; GCL, ganglion cell layer; L, lens; D, dorsal; V, ventral. Scale bar 50 μm.

**Figure S9.**
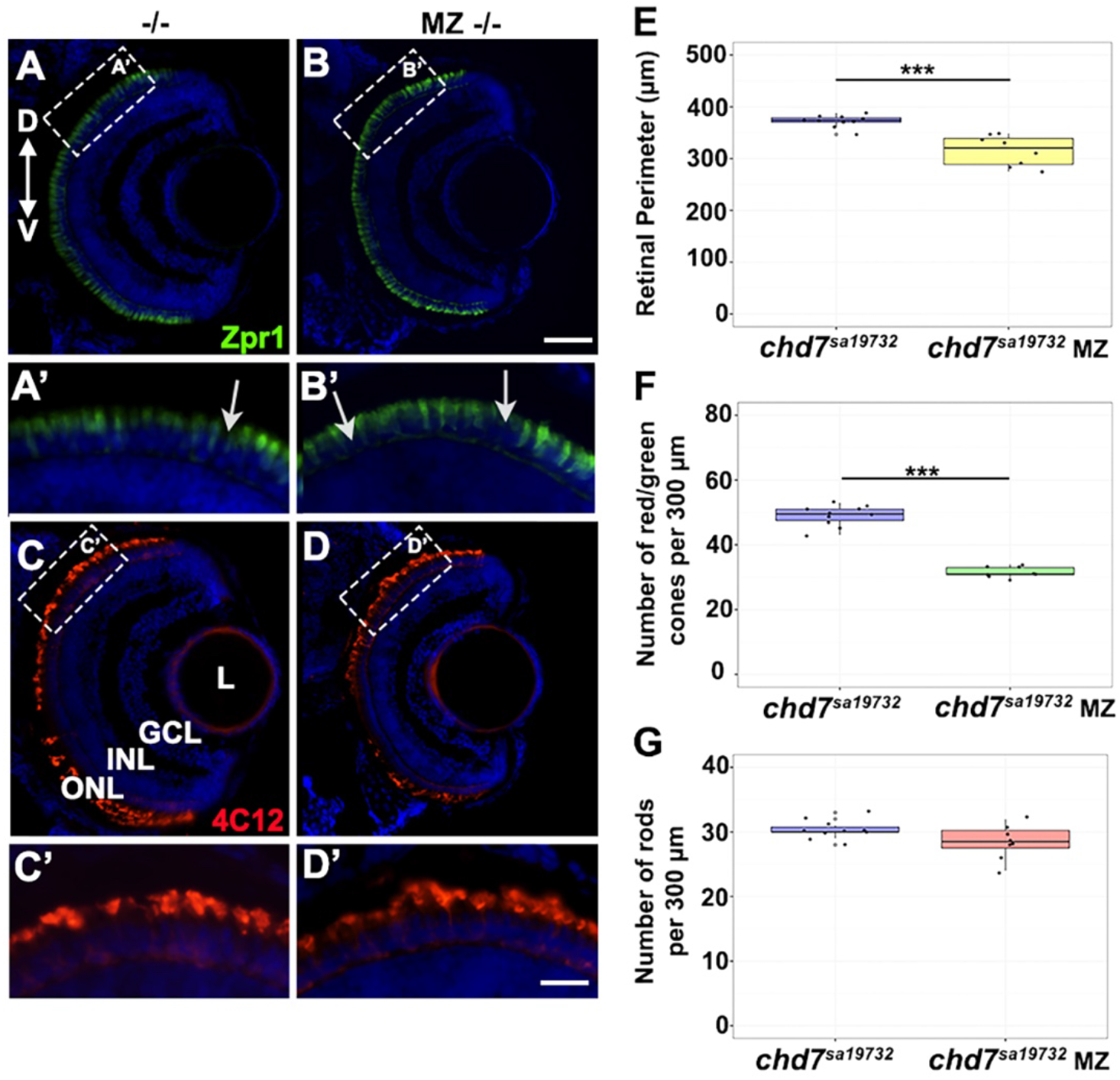
*chd7^sa19732^* maternal zygotic mutant zebrafish display more severe phenotypes compared to zygotic mutants. (A-B) Immunohistochemistry with a red-green cone antibody (Zpr1) in zygotic *chd7^-/-^* (A,A’), maternal zygotic (MZ) *chd7^-/-^* retinas at 5 dpf (B,B’). (C-D) Immunohistochemistry with a rhodopsin antibody (4C12) in zygotic *chd7^-/-^* (C,C’), maternal zygotic (MZ) *chd7^-/-^* retinas at 5 dpf (D,D’). (E) Maternal zygotic mutants have a smaller retina perimeter compared to zygotic mutants. Measurements taken from perimeter of retina normalized to nose to otolith length. (F) Quantification showed decrease in red/green cones in maternal zygotic *chd7^-/-^* compared to zygotic *chd7^-/-^* larvae (number of cones per 300 uM; n=10; ANOVA followed by t-test \*\*\**p* <.001). (G) Quantification showed no difference in rod photoreceptors between zygotic *chd7^-/-^* and maternal zygotic (MZ) *chd7^-/-^* retinas (number of rods per 300 uM). ONL, outer nuclear layer; INL, Inner nuclear layer; GCL, ganglion cell layer; L, lens; D, dorsal; V, ventral. Scale bars 25 μm in A’-D’, and 50 μm in A-D.

**Supplementary Table 1.**
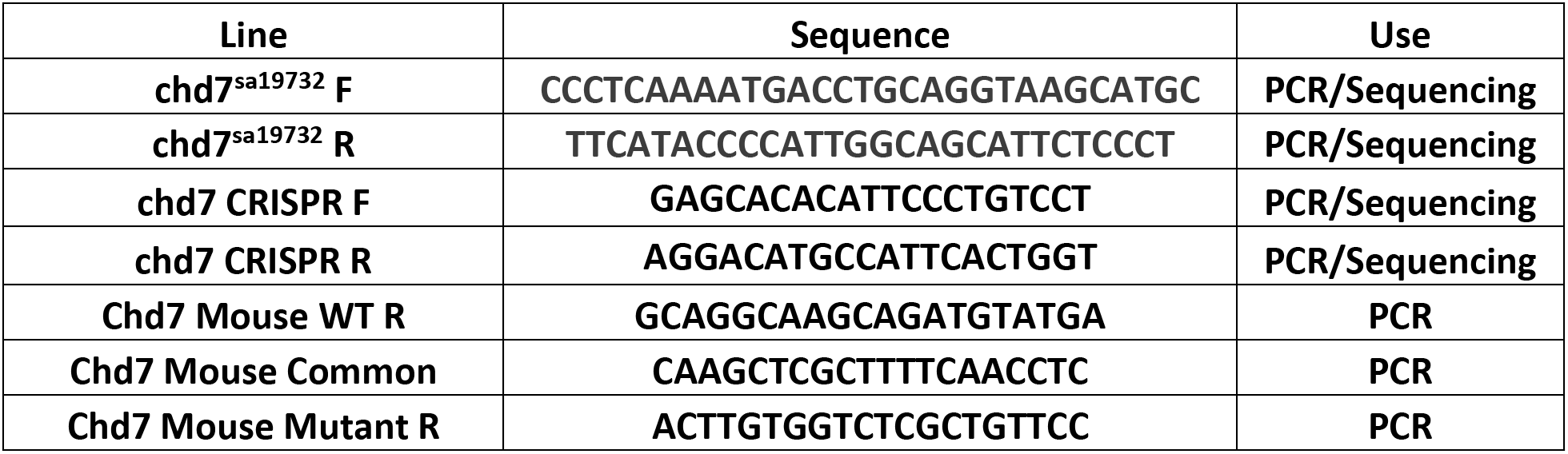
PCR and Sanger Sequencing primers used in this study.

**Supplementary Table 2.** Cellular expression data for *CHD7* from PLatform for Analysis of scEiad (Single Cell Eye in a Disk)

| Predicted Cell Type | Cells # Detected | Total Cells | % | log2(counts+1) |
| --- | --- | --- | --- | --- |
| **Cone** | 3226 | 7854 | 41.07 | 0.49 |
| **Late RPC** | 755 | 1762 | 42.85 | 0.43 |
| **RPC** | 11834 | 31803 | 37.21 | 0.43 |
| **Neurogenic Cell** | 1438 | 3300 | 43.58 | 0.4 |
| **Retinal Ganglion Cell** | 7338 | 19815 | 37.03 | 0.4 |
| **Photoreceptor Precursor** | 948 | 2621 | 36.17 | 0.37 |
| **Rod** | 23340 | 80847 | 28.87 | 0.34 |
| **Corneal Nerve** | 104 | 376 | 27.66 | 0.32 |
| **Muller Glia** | 9719 | 35813 | 27.14 | 0.31 |
| **Blood Vessel** | 2331 | 9744 | 23.92 | 0.29 |

